# Nonlinear sensory integration drives configural olfactory representations

**DOI:** 10.64898/2026.10.09.758135

**Authors:** Eduard Bokman, Ido Padro Kalij, Yael Waxman, Ben Towler, Yuval Balshayi, Yifat Eliezer, Alon Zaslaver

## Abstract

Natural environments are composed of complex stimulus mixtures that can be perceived either as combinations of known components (elemental coding) or acquire entirely new qualities (configural coding). While configural representation is traditionally viewed as a high-order emergent property of cortical processing, the underlying circuit mechanisms remain elusive. Employing *C. elegans*, we demonstrate that the neural foundations of configural representation can be established already at the sensory periphery. For most mixtures, single neuron responses arise through linear integration of the individual components. However, for a subset of mixtures, a small number of sensory neurons exhibit strong nonlinear integration, including NAND/XOR-like logic-gates computations. These nonlinear integrations collectively generate configural population codes. We present a novel quantitative predictor of configural representation and demonstrate that mixtures classified as configural induce emergent synergistic attraction and may lead to violations of rational choice behavior. Together, our findings identify single-neuron nonlinearities as a mechanistic link between sensory circuit dynamics and complex perceptual and decision-making outcomes.

## Introduction

In natural environments, chemical cues rarely appear in isolation, but rather as complex mixtures of numerous components. Animals, including humans, rely on their chemosensory systems to detect and interpret these blends to navigate their environment, forage, find mates, and avoid dangers. Consequently, chemosensory systems face the significant challenge of extracting pertinent information from this chemical complexity to assign meaning to stimuli and drive adaptive behaviors.

Odor mixtures are increasingly discussed in terms of “odor objects”: stable percepts of complex chemical cues, often tied to salient ecological contexts, analogous to object perception in vision ^1–3^. While the analogy and underlying mechanisms remain debated, evidence supports some form of “objectness” across levels of olfactory processing, from early receptor interactions to cortical integration and subjective experience ^4–9^. This framing highlights a trade-off between precision and efficiency: decomposing a mixture into its components preserves information but can be costly and often unnecessary, whereas chunking co-occurring cues into ecologically meaningful entities can speed up decisions while sacrificing details about the parts. Consistent with this, human perception allows mixtures to be processed as individual components (elemental/analytic) or as an integrated object, giving rise to emergent perceptual properties (configural/synthetic), and these modes can coexist within the same stimulus ^1,10–13^. Importantly, configural perception does not necessarily require complete replacement of the component qualities; mixture-specific and component-related qualities may coexist within the same percept ^1^. The balance between these processing modes depends on factors including stimulus identity, component ratio, and mixture complexity ^14–19^.

In vertebrates and insects, mixture processing begins in the periphery: olfactory receptors on olfactory sensory neurons (OSN) are broadly tuned, encoding odor identity combinatorially by patterns of activated receptors/neurons ^20–22^. At this stage, components can interact at the receptor level, producing suppression, partial additivity, or enhancement that reshapes the effective input and can bias perception of blends toward unified “odor objects.” OSN axons then converge onto glomeruli, where lateral inhibition contributes to the sparse mixture representations and has been proposed to support configural effects, though evidence is mixed as to whether early circuits remain largely elemental or shift toward configural coding as complexity increases ^23–25^. Downstream, higher-order regions (piriform and orbitofrontal cortex and amygdala in mammals and mushroom body in insects) further transform these representations to support mixture discrimination and to encode valence and subjective perceptual identity, consistent with increasingly complex perceptual representations along the hierarchy ^3,4,13,26^.

An emerging consensus suggests that configural coding requires complex circuitry dynamics across a multi-layered brain organization. However, the neural mechanisms and computational principles governing configural perception remain poorly understood. To elucidate these principles, it may be advantageous to focus on a simple and well-defined nervous system. To this end, the *C. elegans* neural network offers a powerful platform to address such fundamental questions. Its fully mapped connectome comprises 302 neurons ^27–29^, and advanced imaging techniques allow quantification of whole-network activity dynamics with cellular resolution ^30–32^.

The chemosensory system of *C. elegans* comprises 12 pairs of neurons arranged in a left-right bilateral symmetry ^27,33^. While the number of chemosensory neurons is small, the receptor repertoire is considerably larger, spanning nearly 1,000 olfactory GPCRs, and individual neurons express on average several dozens of these receptors ^34,35^. Accordingly, the sensory system is not strictly segregated into olfactory and gustatory modules. Instead, these neurons respond to both volatile and soluble stimuli: a subset act as direct primary sensors for a target cue while others are recruited to respond via feedback and recurrent synaptic and extrasynaptic signaling ^36–39^.

Here we leverage whole-system functional imaging to investigate how complex chemosensory stimuli are processed in the compact *C. elegans* sensory system. We show that, similarly to vertebrate and insect sensory coding, responses of the *C. elegans* chemosensory system lie on a low-dimensional manifold in neural activity space that tracks perceptual properties like stimulus identity and valence. Within this space, we use the elemental–configural framework to quantify whether mixture representations preserve the component-defined organization or acquire additional mixture-specific structure. Across mixtures, the responses of most neurons are well approximated by a simple linear combination of the responses to the individual cues within the mixture. However, some neurons perform nonlinear integration, including an apparent NAND logic-like gate. Interestingly, these nonlinear responses of individual neurons drive the configural coding of mixtures whereas elemental coding is associated with linear integration. These findings demonstrate that configural representation can arise at the earliest stages of sensory processing, driven in part by nonlinear response dynamics of individual sensory neurons.

## Results

To study how the compact chemosensory system codes mixed stimuli, we measured neural responses to a variety of chemical stimuli presented either individually or as a combination of two or three stimuli (**Figure 1A and Supplementary Figure 1**). Stimuli spanned both valence (attractive and repulsive) and modality (soluble and volatile). To record responses of the chemosensory system, we used transgenic animals expressing the genetically-encoded calcium indicator GCaMP in all amphid sensory neurons ^39–41^. Individual animals were inserted into a custom-made microfluidic device ^42^, and imaged following stimulus presentation (an ON step) and removal (OFF step). We used a fast-scanning confocal system to simultaneously image all chemosensory neurons (two head volumes per second, each with 30–40 slices at a 0.6–0.7 μm z-resolution), providing the necessary spatiotemporal resolution to reliably extract activity from virtually all individual amphid neurons of both lateral sides (22 neurons in total) with the exception of AFDL/R which were often below detection levels. Neuronal responses were calculated as the maximum amplitude in the 7 seconds after a stimulus step^48^.

**Figure 1.**
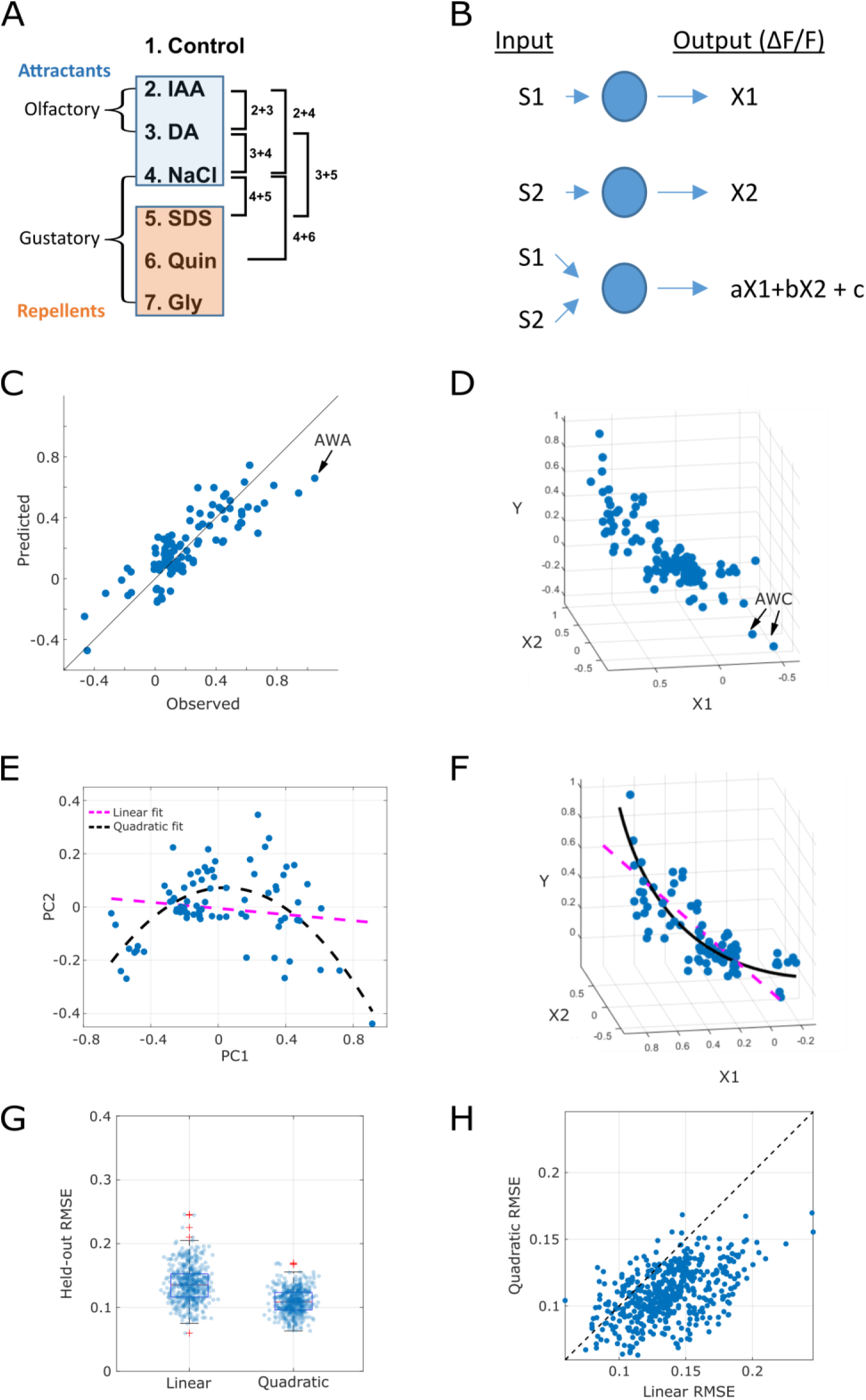
Sensory neurons integrate stimuli combinations in linear and nonlinear fashions. **(A)** The individual stimuli and combinations used in this study. The individual cues consisted of olfactory and gustatory cues, some of which were attractants and some repellents. IAA, Isoamyl alcohol; DA, Diacetyl; NaCl, Sodium Chloride; SDS, Sodium dodecyl sulfate; Quin, Quinine; Gly, Glycerol. **(B)** A linear fit model which postulates that individual neurons simply integrate the responses to the individual stimuli components via a weighted sum. X1 and X2 are the responses of a neuron to the individual stimulus components 1 and 2, and y is the response to their mixture y = aX1 + bX2 + c, where c is the fit intercept. Only neurons that responded above a threshold of 0.1 ΔF/F0 to at least one of the stimuli in the triplet were used for the analysis. The AWC neurons were excluded as outliers. **(C)** Observed *vs* predicted responses. Black line denotes the (1, 1) line. Dots represent all neuronal responses (ON and OFF) to stimuli mixtures with the threshold exceptions specified in B. Combining all neural responses shows a fair fit to the linear model y = aX1 + bX2 + c (R^2^ = 0.65), with a slightly higher weight (b) given to the stimulus eliciting a stronger response (X2). a = 0.39 (p < 0.001) and b = 0.63 (p < 0.001), c = ∼0 (p > 0.05). Arrow points to an example AWA neuron, which is systematically underestimated by the model. **(D)** Scatter plot of neuron responses (ΔF/F0) to combination (Y) against the component responses (X1, X2). For consistency, X1 and X2 were sorted such that X2 was always the component with the strongest response magnitude (in absolute value). The AWC neurons (arrows) were the only neurons who responded with decreased calcium levels, and hence were removed from the fit analysis. **(E)** Projection of the activities onto the principal plane of the parabola. Purple line: linear fit, black line: quadratic fit. Linear RMSE = 0.142, Quadratic RMSE = 0.115 **(F)** Projection of the fit lines back onto the original 3D space. Purple line: linear fit, black line: quadratic fit. **(G)** Distribution of cross-validation RMSE values for the linear and quadratic models folds. Each point represents a single neuron in a single combination. 100 repeats of 5-fold cross-validation with different random seeds. Red line: median, boxes: interquartile range, whiskers: extreme values, red pluses: outliers (>1.5 IQR). **(H)** Fold-by-fold linear vs quadratic RMSE. Points below the diagonal indicate folds in which the quadratic model performed better (87.8%)

Because linear integration is common in olfactory circuits ^43,44^, we first asked whether mixture responses could be explained by a simple linear combination of the responses to the individual components (**Figure 1B**). For each neuron, we fit a multiple linear model of the form *aX*1 + *bX*2 + *c* = *y*, where *X*1 and *X*2 denote the weaker and stronger (respectively) responses of a neuron to the individual components, and *y* is the response to the mixture. This linear model yielded a fairly good fit (*R*^2^ = 0.65), with no intercept, and a slightly larger weight given to the stronger response (*a* = 0.39, *b* = 0.63, *c* ≈ 0), indicating that integration of stimulus combinations by individual sensory neurons is, overall, well explained by a linear function of the responses to the individual stimuli when presented alone (**Figure 1C**).

A subset of neurons, however, showed strong and consistent deviations from the linear model (**Supplementary Figure 2**). For example, in AWA neurons, the linear model systematically underestimated mixture responses, consistent with nonlinear integration (**Figure 1C, Supplementary Figure 2**). Other neurons (*e.g.*, ASH, ASER, ASJ) deviated from the linear model in response to specific stimulus combinations. Plotting mixture responses against the individual responses revealed a parabolic-like structure (**Figure 1D**). We therefore fitted a parametric function to the data by projecting the points onto the principal plane of the parabola and then fitted a quadratic equation (**Figure 1E-H**, and see methods). The quadratic model provided a better fit to the data than the linear function with lower, and more consistent prediction error (5-fold cross-validation; quadratic vs linear mean RMSE: 0.109 vs 0.137, std: 0.019 vs 0.028, **Figure 1G,H**). Bayesian information criterion (BIC) also favored the quadratic model over the linear model (-351.17 vs -318.22; ΔBIC = 32.944, **Supplementary Table 1**). Overall, these results indicate a sizable nonlinear component in the integration of complex stimuli in the chemosensory neurons.

### Chemosensory neurons employ nonlinear integration of competing stimuli

To better understand these potentially nonlinear integration processes, we considered the functional consequences of stimuli integration. In this context, sensory neurons typically exhibit several possible integration motifs: (i) Overshadowing, where the response of a neuron to the mixture resembles the response to one of the components; (ii) Suppression, or weighted average; (iii) Synergy, or positive weighted sum; and (iv) Inhibition, or negative weighted sum. Overshadowing and suppression can arise from linear integration, whereas synergy can reflect either linear additivity or nonlinear amplification. Inhibition relies solely on nonlinear integration.

We found instances of all these motifs across chemosensory neurons (**Figure 2A-F**). Among the responding neurons, the most common form of integration was overshadowing, accounting for ∼53% of all cases, and suppression, which accounted for an additional 25%. Thus, purely linear integration was observed in over 75% of the cases. Synergy and inhibition accounted for 18% and 4%, respectively (**Supplementary Figure 3**).

**Figure 2.**
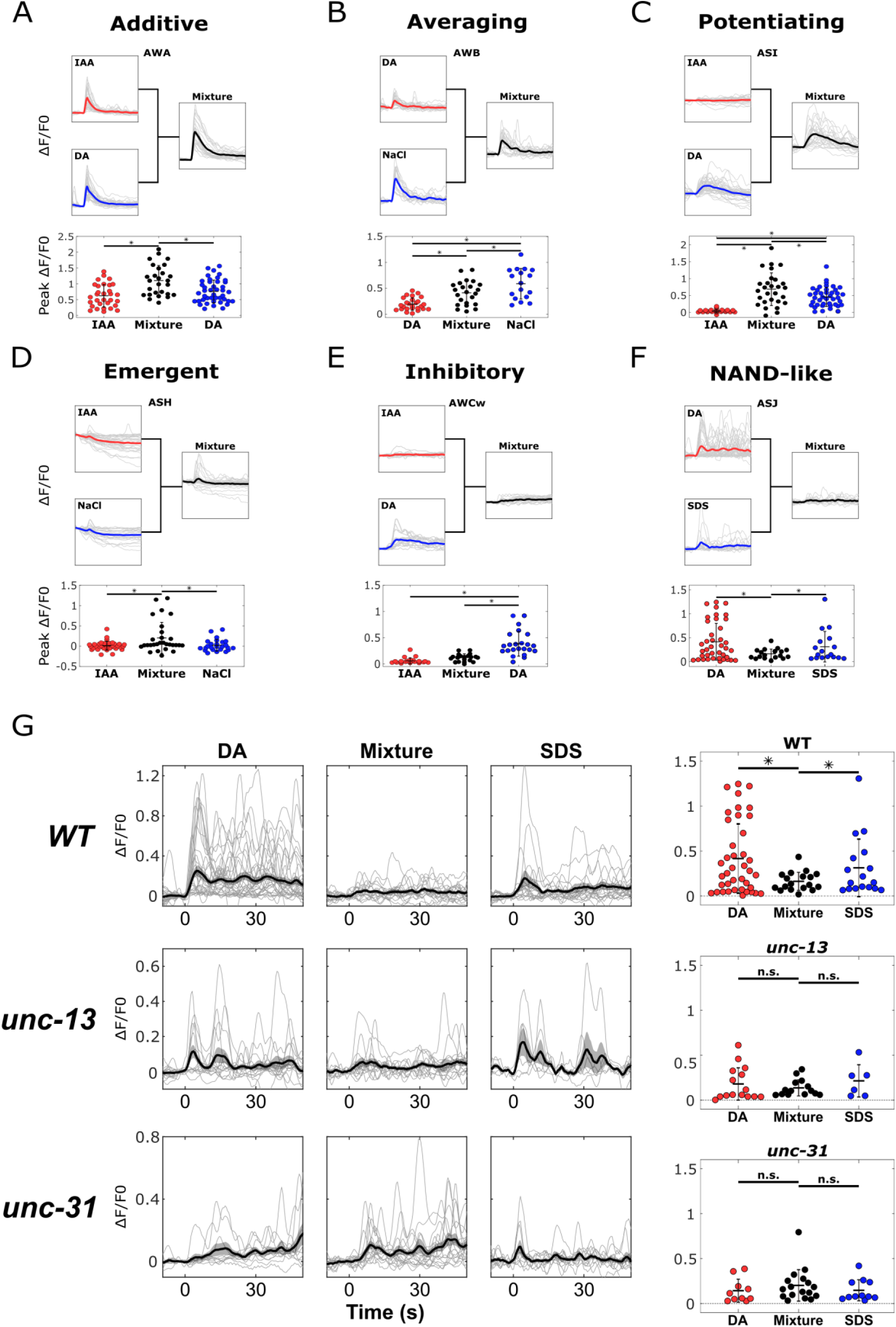
Sensory neurons employ a variety of linear and nonlinear integration strategies. **(A-F)** Examples of integration strategies observed in individual sensory neurons. (A) an additive effect between IAA and DA in the AWA neuron. (B) An averaging effect in response to the combination of NaCl and DA in the AWB neurons. (C) A potentiating effect in response to the combination of IAA and DA in the ASI neuron. (D) An emergent effect in response to the mixture of IAA and NaCl in the ASH neuron. (E) An inhibitory effect in the AWC neuron in response to the combination of IAA+DA. (F) A NAND/XOR-like logic gate in the ASJ neurons in response to combination of SDS and DA. Note that while the ASJ neurons respond to each of the stimuli when presented alone, they do not respond to their combination. Top: Single neuron traces (grey) and population mean traces of the responses. Red and blue: individual stimulus components; black: their mixture. Bottom: Peak response amplitudes (ΔF/F0). * p<0.05. **(G)** The NAND/XOR logic-like gate is the outcome of the network activity rather than a cell-autonomous effect. Responses of the ASJ neurons to DA, SDS, and their mixture. Left: Individual ASJ traces (grey) and mean trace (black). Shaded area indicates standard error of the mean. Right: peak amplitudes. * p<0.05.

We also observed more nuanced examples of some of these effects that allow for a more granular classification. For example, we observed positive overshadowing (*A* > 0, *B* = 0, *AB* = *A*) in the AWB neurons in response to the mixture DA+NaCl (**Figure 2B**), and negative overshadowing (*A* > 0, *B* = 0, *AB* = 0) in the AWC neurons in response to DA+IAA (**Figure 2E**). Synergy could be further subdivided into additive (*A* > 0, *B* > 0, *AB* > *max*(*A*, *B*), e.g. AWA in DA+IAA, **Figure 2A**), potentiating (*A* > 0, *B* = 0, *AB* > *A*, e.g. ASI in DA+IAA, **Figure 2C**), and emergent (*A* = 0, *B* = 0, *AB* > 0, e.g. ASH in IAA+NaCl, **Figure 2D**). Finally, a striking case of inhibition was found in the ASJ neurons in response to DA+SDS: This neuron type responded to both individual stimuli but not to the mixture, reminiscent of a NAND or XOR gate (*A* > 0, *B* > 0, *AB* = 0, **Figure 2F**).

Effects such as additive synergy and overshadowing could arise cell-autonomously, for example, due to interactions at the olfactory receptors, or within downstream intracellular signaling pathways. Other effects, like emergent synergy, could stem from cell-autonomous threshold effects. However, the integration mechanism underlying the extreme case of inhibition is less obvious: How can the ASJ neurons respond to DA and SDS when presented separately, yet fail to respond when presented as a mixture? To delineate whether this is a cell-autonomous or a network-mediated effect, we employed two mutant strains, each defective in one network communication pathway: *unc-13*, which is impaired in synaptic transmission, and *unc-31*, which is defective in neuropeptide release. Although ASJ responses were slightly attenuated in the mutant strains, in both *unc-31* and *unc-13* mutants, the mixture responses were comparable to the component responses, suggesting that the inhibition is not cell-autonomous but rather results from network feedback effects (**Figure 2G**).

Together, these findings indicate that the majority of the integration processes follow a simple linear weighted sum model, while a wide array of nonlinear interactions in some neurons add complexity to the representation of odorant mixtures.

### Sensory representations lie on a low-dimensional nonlinear manifold in neuronal space

Thus far, we considered activity dynamics of each neuron separately and irrespective of the activity of other neurons. However, perceptual representation of a stimulus is an emergent property of the coordinated activity of the collective neuron ensemble. To examine how mixtures are represented in the compact chemosensory system, we constructed a sensory representation space by performing linear (PCA) and nonlinear (Isomap) dimensionality reduction on the neural activities (**Figure 3A and Supplementary Figure 4A-B, Supplementary Note 1**). The combined responses showed a robust low-dimensional structure, with the nonlinear manifold found by Isomap capturing 87% of the variance in only 3 dimensions, while the linear PCA captured 43% (though both methods provided good visual separation of the stimulus clusters). Including the mixture responses in the analysis only slightly reduced the variance explained by the nonlinear manifold to 79%, suggesting that the individual component manifold captures the mixtures as well (**Supplementary Figure 4I**).

**Figure 3.**
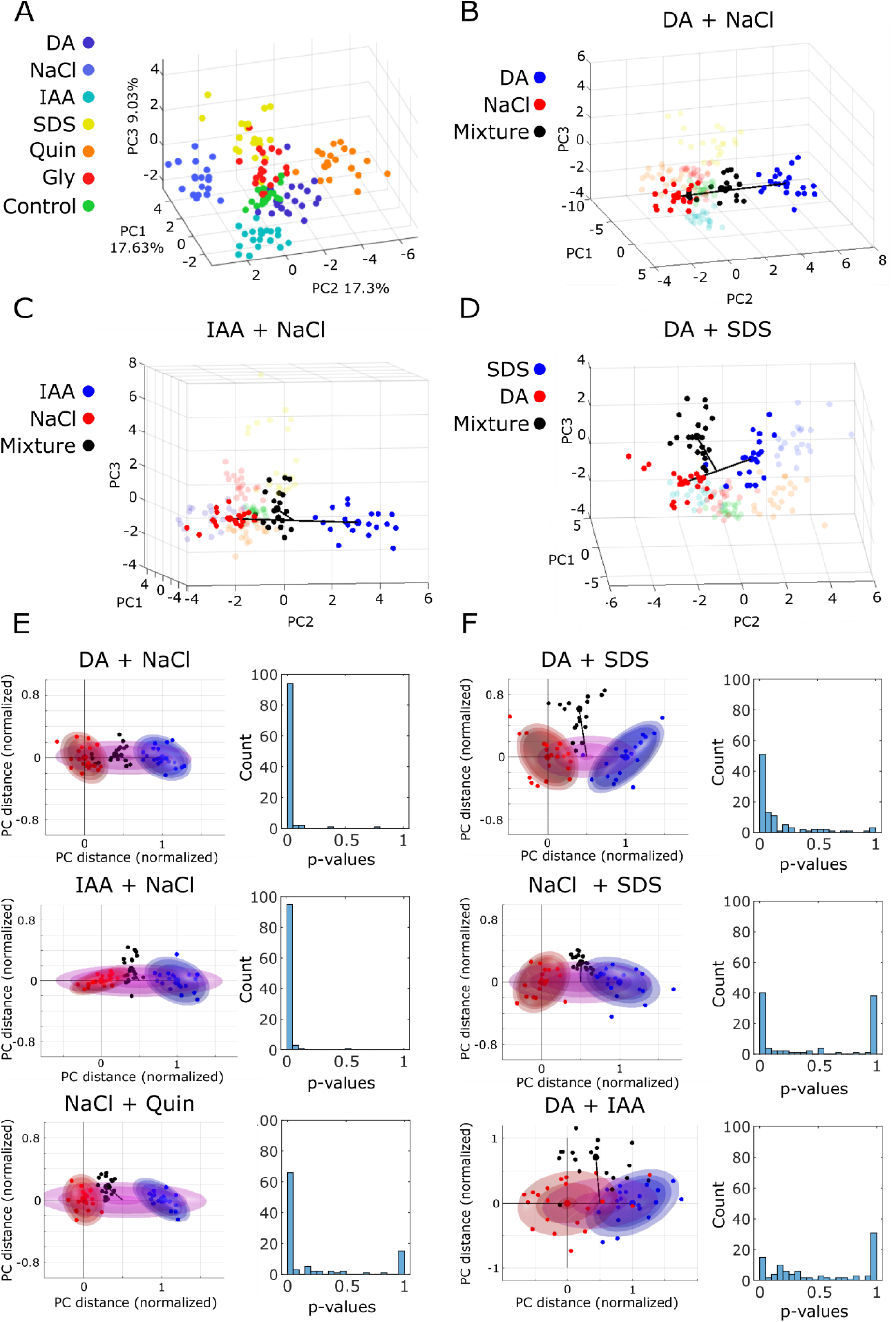
Sensory representations lie on a low dimensional manifold, some of which occupy novel distinct niches in the neuronal space. **(A)** Scatter plots of the responses to individual stimuli in the first three components of the PC space. Each point in the space represents an individual worm, described by 13 ON and 13 OFF responses (see Methods). **(B-D)** Examples of responses to mixtures in the PC space in relation to their components showing no deviation from the centerline (B), weak deviation (C), and strong deviation (D). **(E-F)** Left: Mixture and component clusters in PC space, rotated to align the centerline with the x-axis. Points represent individual worms in the components (red and blue) and the mixture (black). Concentric spheres represent the probability density function of the Gaussian mixture model fit to each component separately (red and blue), and to the combined cluster that includes all worms from both components (purple). The values of the PDF, denoted by the shaded regions in the spheres, are consistent within each panel. Distances were scaled to make the centerline be of length 1. Right: Distribution of p-values obtained by a paired Wilcoxon signed-rank test between the PDF value of the mixture points in the combined cluster and the larger PDF value of the mixture in either of the individual clusters. Small p-values (p<0.05, first column) indicate a significantly higher probability that the mixture belongs to the combined cluster, placing it on the centerline (consistent with elemental processing). The p-values were obtained by 100 random resamplings of 20 worms per condition.

### Mixed stimuli occupy novel niches in the representational space

The ensemble analysis showed that each stimulus occupies a well-defined region of a low-dimensional “perceptual space” (**Figure 3A**). Each point in this space is a single worm, represented as a vector of length 26 containing the responses of the 13 neurons to stimulus presentation and removal. Mirroring similar dimensionality reduction approaches in neuronal, chemical, and perceptual studies of olfactory stimuli, the *C. elegans* neuronal space reflects both odor similarity and higher-order features such as valence (**Supplementary Figure 4C-G and Supplementary Note 1)**. For example, the attractive, food-associated odors DA and IAA cluster closer to each other than to the aversive stimulus SDS, suggesting similarity of their representations. Thus, projecting mixtures onto this space could provide insights into their perceptual quality.

Mixture processing ranges from elemental, in which the mixture is represented through its component-related qualities, to configural, in which the blend acquires a novel quality that may either coexist with or replace the qualities of the individual components ^1^. In the low-dimensional neural manifold, a perfectly elemental representation, where both components contribute equally, should place the mixture cluster directly on the line connecting the two component clusters. Displacement away from this line toward a region not occupied by either component would indicate a representational shift consistent with configural coding (for example **Figure 3B**). Additionally, the distances of the mixture cluster to each of the component clusters should reflect component dominance: in the extreme case, where the mixture completely overlaps one of the components, the presence of the second component is not expressed in the neuronal activity and is unlikely to contribute to perception. Across all combinations we assayed, mixture clusters fell between the corresponding component clusters, with the distance from the mixture to each component being smaller than the distance between the two components (**Figure 3B**, and **Supplementary Figure 5**). This is consistent with the observation that neurons mostly integrate mixtures using weighted averages and suggests that mixture representation exists on a continuous axis between the individual components.

Notably, some mixtures deviated from this axis (**Figure 3C-D**). Such deviations are of particular interest because they suggest that mixture representations can occupy a representational niche that is different from the sum of their parts. To quantify the significance of this deviation, we fitted Gaussian mixture models to each component cluster as well as to a pooled cluster that contained all the points from both components (**Figure 3E-F**). We then compared, for each mixture point, the probability density under the two component models versus the pooled model. If the mixture lies near the line connecting the single components, then the value of the probability density function (PDF) of the pooled model is expected to be significantly larger than those of the individual models. By contrast, if the mixture deviates from the centerline, then the PDF value of the pooled model should be equal to, or lower, than the PDF of the individual models. We found that mixtures lie on a spectrum of deviation from the central axis: Some mixtures, such as DA+NaCl and IAA+NaCl fall directly in the middle of the component clusters (**Figure 3E**), consistent with linear weighted mean integration, and elemental representation. Other mixtures, like NaCl+Quin, DA+SDS, and NaCl+SDS, deviate from the centerline (**Figure 3F**), shifting into a novel region of the space, suggesting the possible emergence of a configural representation. Finally, DA+IAA fell well outside the component regions, indicating a pronounced representational shift and strong configurality.

Taken together, these results indicate that *C. elegans* encodes most binary odor blends as elemental, linear interpolations that position mixture representations along the continuum between their component clusters. Nevertheless, systematic off-axis displacements for specific blends suggest that the sensory manifold can also support emergent configural codes, generating novel representational qualities not predictable from their parts.

### Individual neurons showing nonlinear integration contribute to configurality

We next studied the relationship between configurality and nonlinearities in the sensory neurons using a synthetic knockout-and-rescue analysis (**Figure 4A-E**). We first constructed a null model of a perfectly elemental representation by averaging randomly sampled pairs of worms taken from the components of a mixture. When projected onto the PC space, this results in a cluster of points scattered around the center point between the component clusters. We then re-inserted the empirically measured mixture responses one neuron at a time, projected the new points onto the PC space, and quantified the deviation of the resulting points from the midpoint. We developed the following approach to quantify this deviation (**Figure 4A**): we defined the line connecting the centers of the component clusters as the major axis of an ellipse (*a*). This axis, together with the center of the reconstructed mixture cluster, defines a unique ellipse with a minor axis *b*. Configurality was taken to be the ratio 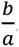 In a perfectly elemental representation, the mixture falls directly on the line, giving *b* ≈ 0, and Configurality 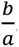 = 0. This ratio increases as the mixture drifts away from the connecting line *a*, while also accounting for the distance between the component clusters, requiring a larger deviation the further the components are from each other. Interestingly, we found that the deviation was larger when component clusters were closer to each other, not only relative to their distance, but also in absolute terms, suggesting that configurality increases with the similarity of the components (**Supplementary Figure 6**). Importantly, this quantitative approach to configurality strongly agrees with our previous statistical definition of configurality (compare results in **Figure 3E-F and Figure 4B**).

**Figure 4.**
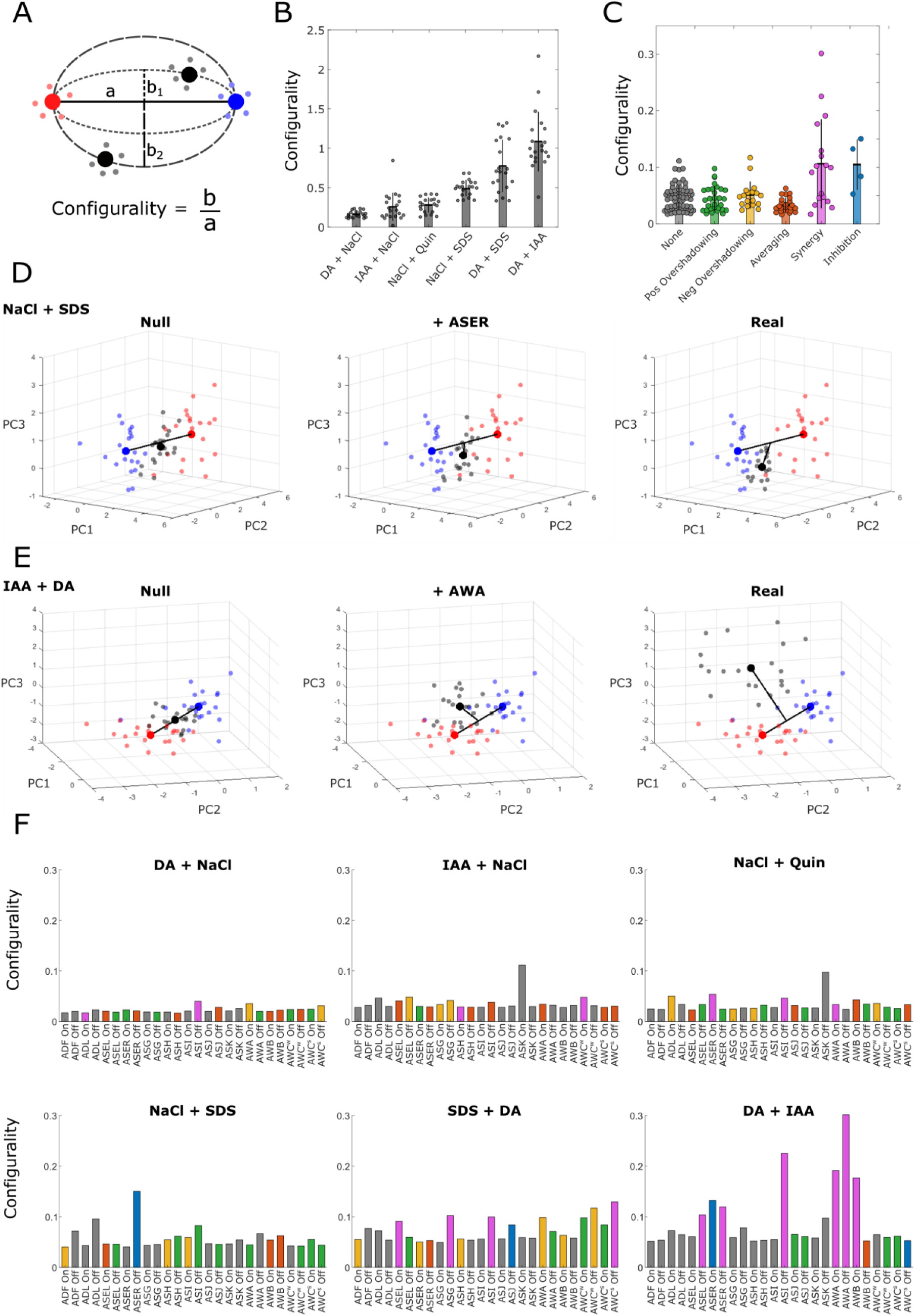
Nonlinear integration in individual neurons contributes to configurality. **(A)** Diagram of configurality measure. Single worms (small circles) and cluster centroids (large circles) of individual mixture components (red, blue) and the mixture itself (black) in PC space. The line connecting the centers of the component clusters is denoted (a). The centerline and mixture cluster define a unique ellipse with major axis (a) and minor axis (b). Configurality is defined as the ratio b/a, with small deviations giving flatter ellipses (b1) and resulting in low configurality, and stronger deviations giving larger ellipses (b2) and stronger configurality. **(B)** Configurality of the tested mixtures. Points represent overall configurality of the mixture cluster in a random resampling of worms. Bars represent the mean configurality of all resamplings. Error bars denote standard error of the mean. **(C)** Contribution of an individual neuron to the configurality of a mixture cluster, grouped by integration strategy. **(D-E)** Examples for contribution to configurality by a single neuron integrating by inhibition (D) and synergy (E). Component clusters are in red and blue, mixture cluster in black. Left: Null mixture (mean of components), middle: null mixture with a single real measured neuron, right: full measured mixture. **(F)** Contributions of individual neurons to configurality. Colors correspond to the integration strategies in (C). Green, positive overshadowing; Yellow, negative overshadowing; Orange, suppression; Pink, synergy; Blue, inhibition.

Grouping the neurons by integration strategies revealed that the canonically nonlinear integrations (synergy and inhibition) contributed more strongly to configurality than the linear ones (overshadowing, suppression) (**Figure 4C**). For example, the inhibitory integration in ASER in the NaCl+SDS mixture (**Figure 4D**), and the synergistic integration in AWA in DA+IAA (**Figure 4E**) were sufficient to shift the reconstructed mixture cluster a sizable portion of the way toward the empirically observed mixture cluster. Nonlinear motifs were also more frequent in the three most configural mixtures than in the more elemental ones (**Figure 4F and Supplementary Figure 3**), and their occurrence correlated strongly with configurality (**Supplementary Figure 7**). For example, the strongly configural DA+IAA mixture (**Figure 4B**) showed considerable contributions from multiple neurons exhibiting pronounced synergy or inhibition (**Figure 4C, F**).

Taken together, these results suggest that configural representations are driven by nonlinear integration within chemosensory neurons. It should be noted that this is a conservative approach, as it relies on the contribution of individual neurons to the first few components of the PC space. Consequently, neurons that exhibit strong nonlinear integration but contribute little to the first PCs (for example, ASJ inhibition in DA+SDS; **Figures 2F and 4F**) may be underweighted.

### Configural representation in neuronal space predicts behavioral outcomes

We next asked whether divergence in mixture representations across the neural ensemble is reflected in behavior. To test this, we performed chemotaxis assays to binary mixtures and their individual components (**Figure 5A**). We chose the DA+IAA and DA+NaCl mixtures as the most extreme cases of configural and elemental representations, respectively. Because all components are attractive, purely elemental integration should result in additive attraction, whereas a novel configural quality could alter the perceived attractiveness of the components and result in antagonistic or synergistic integration. First, we constructed dose-response curves for each individual component and fit a sigmoidal hill function to each (**Figure 5B**). We then measured chemotaxis to mixtures across a range of component concentrations, chosen to span the range of attractiveness (0-1 chemotaxis index). To look for synergy or antagonism, we tested dose equivalence for each mixture by calculating the Loewe’s additivity index (LI), an index commonly used to detect synergism in drug combinations ^45,46^. Loewe’s index asks whether the two components in a mixture behave as interchangeable contributors to the same behavioral effect. A value of 1 indicates that the fractional contributions of the components are fully additive. If the sum is < 1, then lower concentrations of the components are required to achieve the same effect, indicating that the mixture is synergistic. Similarly, a sum > 1 indicates antagonism. For a given mixture dose pair (*x_mix_*, *y_mix_*) that produces effect *CI_mix_*, we used the single-odor calibrations to determine the solo doses *x_solo_*(*CI_mix_*) and *y_solo_*(*CI_mix_*) that would produce the same effect. We then computed the Loewe index as the sum of fractional contributions:

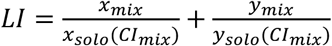

**Figure 5.**
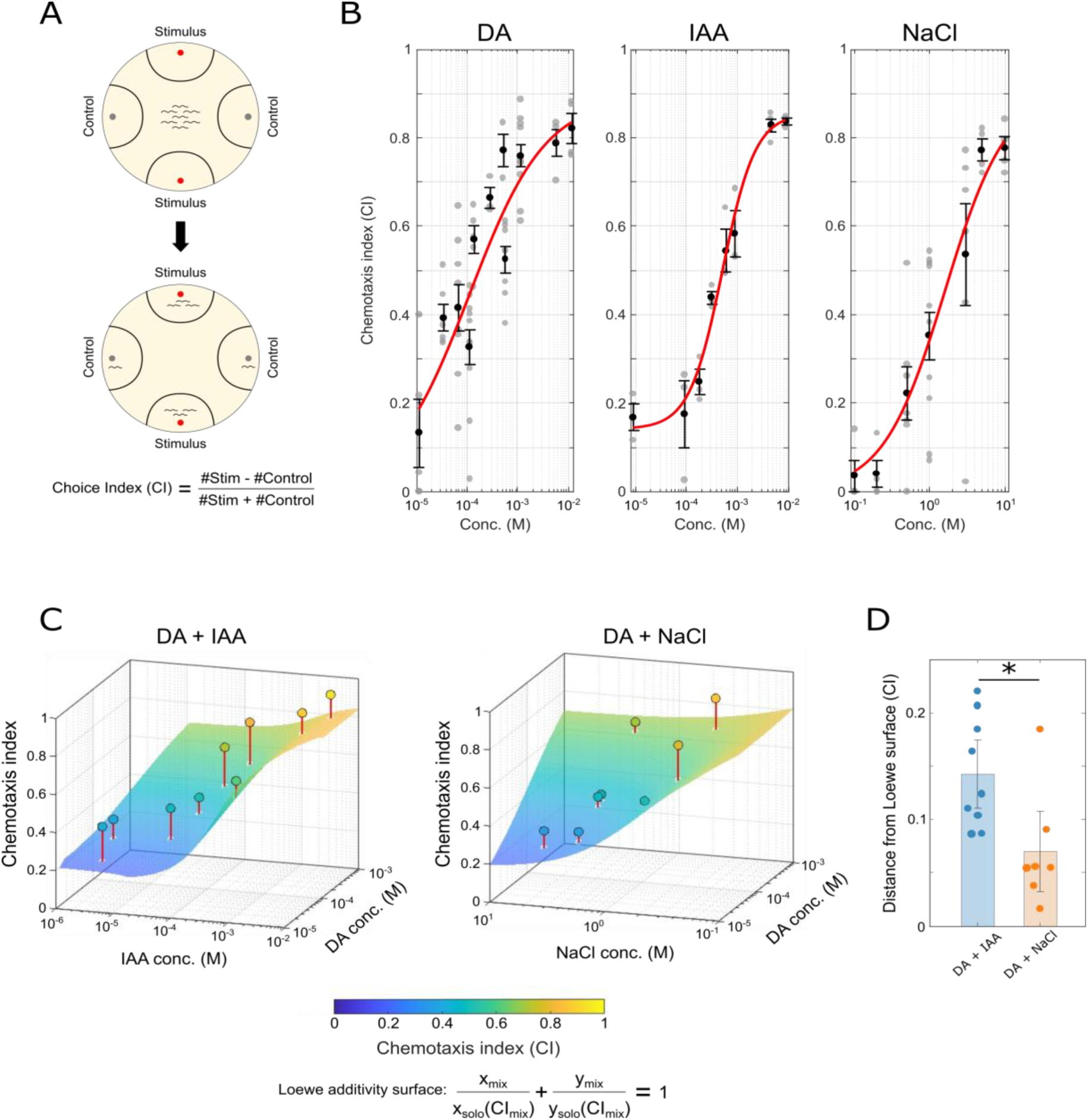
Configural mixture processing is associated with emergent, synergistic behavioral responses. **(A)** Illustration of the choice assay setup. Two points of the stimulus (single component or mixture) diluted in ethanol are placed on opposite ends of a 9 cm plate, and two points of ethanol are placed on the perpendicular ends as controls. Worms are placed in the middle of the plate and allowed to chemotax for 1 hour. Choice index was calculated as # of worms at stimulus minus # of worms at control divided by the total number of worms at the end points. **(B)** Dose-response calibration curves for the individual components. Points represent day averages of three technical repeats per day. Black bars indicate standard error of the mean for all repeats of the same concentration. Red curves represent the fitted logistic model. **(C)** Predicted chemotaxis under Loewe dose additivity based on the calibration curves in (B). The colored surfaces represent the expected Loewe response *CI_L_* for each pair of doses, computed from the isobole equation 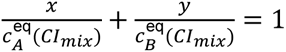 (see methods). Filled circles mark the measured mixture responses; red stems indicate the vertical deviation from the additive prediction surface (positive = synergy, negative = antagonism). **(D)** Summary of deviations from additivity for the configural DA+IAA and elemental DA+NaCl mixtures. This deviation is significantly higher for the configural DA+IAA mixture (* p = 0.017, two-sided t test).

The elemental DA+NaCl mixture was largely additive, with mildly synergistic responses at the high end of the tested concentrations (**Figure 5C-D**), likely stemming from small but cumulative neuronal contributions, or, possibly, real synergy that emerges at effective concentrations higher than were tested in the neuronal recordings. In contrast, the configural DA+IAA mixture maintained a robust synergistic integration across the entire range of tested concentrations, consistent with an emergent attraction to the mixture (**Figure 5C-D**).

Combined, our findings raise the prediction that mixtures of highly similar components should elicit strong nonlinear interactions in individual sensory neurons (**Figure 1, 4F, and Supplementary Figure 6)**, and, consequently, non-additive behavioral outcomes (**Figure 5**). We therefore tested this prediction by analyzing sensory responses and behavioral outputs to the mixture of Isoamyl alcohol and Benzaldehyde, two attractive food-borne odorants that elicit similar sensory responses ^38,47,48^. Our findings support the prediction: individual neurons (ASE, AWC, ASI) showed nonlinear integrations, including emergent and inhibitory effects (**Supplementary Figure 9A-C**), as well as strong behavioral synergy where the attraction to the mixture was significantly stronger than the attraction to a simple additive prediction (**Supplementary Figure 9D-F**).

We next postulated that configural representations may affect rational choice making. For example, the emergence of a novel sensory property in a mixture may lead to violation of the key transitivity paradigm. According to this paradigm, each choice is associated with a utility (value), and rational animals should consistently prefer choices with higher utilities. Elemental mixtures, whose components are processed independently, should preserve these absolute utilities, thus leading to consistent transitive preferences. In contrast, a combination of choices that form a novel configural representation is likely to create a new, context-dependent value, and hence to disrupt the hierarchical utilities associated with offered choices. Such disruption may violate transitivity and, consequently, lead to irrational decision-making. To study this intriguing possibility, we analyzed a comprehensive choice-assay database that scored preference between many different pairs of chemical choices, including the configural DA+IAA mixture, and the elemental DA+NaCl, and IAA+NaCl mixtures ^40^. Strikingly, compared to the elemental pairs of stimuli, the triplets containing the DA+IAA pair were significantly more likely to violate transitivity (**Supplementary Figure 8 and Supplementary Note 2**).

Taken together, these behavioral results provide a direct link between our computationally defined elemental/configural representation framework, driven in part by nonlinear integration in sensory neurons, and the behavioral outcomes elicited by complex stimulus mixtures.

## Discussion

Here, we investigated the neural mechanisms that distinguish elemental from configural processing of odor mixtures. We found that nonlinear integration in individual sensory neurons transforms an elemental coding strategy into a configural representation. Furthermore, we developed a novel metric to quantify and predict configurality based on neural dynamics alone. Subsequent behavioral assays validated these predictions, further supporting our mechanistic findings.

We show that most sensory neurons primarily employ the linear overshadowing and suppression strategies when integrating mixtures (**Figure 1 and Supplementary Figure 3**). This linear integration aligns well with established models of olfactory coding in other organisms, suggesting evolutionary conservation of elemental processing mechanisms. However, the presence of distinct nonlinearities, including synergy and inhibition, within specific neurons introduces complexity to mixture representation. These nonlinear interactions were particularly surprising given their typical association with higher-order processing regions in other organisms ^13,23,25,49,50^. Importantly, these effects are unlikely to stem from physicochemical interactions, as no chemical reactions are known to occur between the components in any of the tested combinations (see methods).

Although nonlinear responses in deeper cortical circuits often arise from recurrent network signaling, those in primary sensory neurons have traditionally been ascribed to receptor-level mechanisms such as saturation or non-competitive binding ^51,52^. We found the pronounced inhibition in ASJ neurons in response to DA+SDS to be network-mediated, rather than cell-autonomous, as evidenced by its dependence on synaptic and extrasynaptic signaling through UNC-13 and UNC-31 (**Figure 2**). While the underlying mechanisms of other nonlinear interactions in the neurons remain unclear, our results suggest that network activity can generate sophisticated computations at the very first stage of sensory processing and underscore the pivotal role of circuit-level signaling in shaping odor-mixture representations throughout the sensory hierarchy.

When analyzed at the ensemble level, sensory responses occupied a low-dimensional nonlinear manifold (**Figure 3 and Supplementary Figure 4**). This manifold divides the response space into distinct regions along both physicochemical and salience axes, as seen in the clear separation of stimuli by identity and valence. Similar manifolds have been consistently reported across multiple studies employing diverse methods and species ^26,53–56^. These span the entire sensory processing cascade from the physicochemical properties of molecules, through cortical recordings and behavioral assays, and up to subjective descriptions of perceptual experiences. These spaces are not equivalent, but their shared relational organization suggests that neural population geometry can preserve stimulus properties relevant to sensation, valuation and perception across species. This correspondence provides a principled basis for applying the elemental-configural framework to *C. elegans*: mixtures positioned primarily along the component axis preserve component-related structure, whereas perpendicular displacement reflects the emergence of mixture-specific structure.

Despite the complexity of neural responses, and in keeping with recently published results ^57^, mixtures generally occupied predictable positions within this perceptual space, typically along an axis defined by their constituent components. A subset of mixtures, however, showed clear displacement from the component axis. Importantly, the interpretation of such displacement depends on the organization of the processing system and the geometry of the neuronal activity space. In labeled-line systems, or systems with more orthogonal processing channels, such as the mammalian olfactory epithelium, linear co-activation of separate channels can place a mixture off the axis connecting its components without implying configural processing. By contrast, the strongly interconnected *C. elegans* sensory system exhibits correlated, low-dimensional population dynamics that share relevant features with downstream mammalian cortical representations rather than with a purely epithelial input layer. This organization, combined with the predominance of weighted-average integration across *C. elegans* sensory neurons, suggests that deviations from the empirically expected component axis are consistent with emergent configural representations.

To quantitatively assess this configurality, we developed novel statistical and analytical methods that measure deviations from the linear axis connecting mixture components (**Figure 4**). This analysis revealed a spectrum of representational shifts ranging from purely elemental to strongly configural, with nonlinear neuronal integration patterns strongly correlating with these configural shifts.

Our synthetic knockout-and-rescue experiments clearly identified individual neurons whose nonlinear responses contributed to configural representation. Specifically, neurons exhibiting synergy and inhibition substantially altered the positioning of mixtures within the ensemble activity manifold. This direct link between neuronal response nonlinearities and configurality demonstrates that complex contextual processing of environmental cues starts already at the sensory layer, possibly aided by top-down feedback signaling.

Configural representations were notably prevalent in mixtures with structurally and ecologically similar components (**Supplementary Figure 6**). This finding aligns with ecological theories suggesting that sensory systems are adapted to integrate stimuli based on shared functional relevance. Indeed, mixtures that deviate significantly from elemental integration, such as DA+IAA, might reflect distinct ecological niches or environmental scenarios demanding unique behavioral responses. Both DA and IAA are prominent attractive molecules emitted by natural *C. elegans* food sources ^58–60^ and detected by the AWA and AWC neurons ^37–39,61,62^. Simultaneous presentation of both chemicals could serve as a high confidence signal for the presence of food and would therefore be integrated as a single cohesive “odor object”. The configurality of the DA+IAA mixture is further supported by behavioral experiments where the mixture elicited a stronger attraction response than predicted by the sum of its parts, even when accounting for the total concentration of the stimulus ^63^. This suggests the emergence of a novel mixture-specific quality, consistent with configural perception.

Nevertheless, chemotaxis does not reveal whether the emergent property reflects odor identity, intensity, valence or their combination. Associative conditioning and generalization between the mixture and its components would help distinguish weak configurality, in which component and mixture-specific qualities coexist, from robust configurality, in which the mixture is processed predominantly as a distinct odor object. Additionally, it is likely that the identity of an odor object is determined not only by the components themselves, but also by their ratio ^18^. While we did observe behavioral synergy in the DA+IAA response in a small range of concentration ratios, we did not systematically examine all possible ratios, and the robustness of the odor object across mixture compositions remains to be determined.

The other potential odor object we observed consisted of diacetyl and SDS. Though SDS is not a naturally occurring organic substance, *C. elegans* might encounter similar compounds in the form of surfactants such as serrawetins and rhamnolipids ^64–66^. Surfactants are produced by some species of pathogenic bacteria and are generally repulsive to the worms. Some of the same bacteria also emit diacetyl. It is possible that the combination of diacetyl and a strong surfactant constitutes a clear signal of potential harm to the worm. If that is the case, it could be beneficial to encode a bouquet of purely attractive odors like DA and IAA differently from the overlapping DA and SDS mixture. Assigning each mixture a distinct odor object that signals either “food” or “danger” is one way to achieve this. Such context-dependent coding is likely advantageous for organisms navigating complex chemical environments.

Importantly, our findings also reveal a continuum between elemental and configural representations, rather than a clear dichotomy. Utilizing the ellipse metric (b/a ratio, **Figure 4A**) to quantify mixture displacement, we observed that the magnitude of displacement scales with the number of neurons exhibiting nonlinear interactions within the ensemble. This indicates a graded spectrum of perceptual coding strategies that reflects the mixture’s composition and ecological relevance. Similar graded phenomena have been reported in psychophysical studies of olfactory perception in rats and humans ^13,67,68^, supporting the idea that perceptual configurality arises as a continuum determined by component similarity and contextual relevance. At the far end of this continuum, highly complex mixtures in humans collapse onto a single, unidentifiable percept dubbed ‘olfactory white’ ^19^. Our data hint at an analogous convergence in *C. elegans* (**Supplementary Figure 4H and Supplementary Note 1**), with the contraction of the perceptual space as the number of components increases. Our observations thereby provide quantitative and mechanistic support for a nuanced model in which elemental and configural processing are not mutually exclusive but coexist and interact dynamically within sensory neural circuits.

The observed neural encoding patterns in *C. elegans* align remarkably well with contemporary theories of configural perception and chunking, a process whereby individual pieces of information are grouped to form a single cohesive object ^69^. The nonlinear combinatorial responses recorded via calcium imaging indicate that odor mixtures activate distinct neuronal assemblies that cannot simply be inferred by summing individual odorant responses. This neural strategy is consistent with the efficient coding hypothesis ^70–72^, where limited sensory resources are optimized by compressing multiple stimulus components into meaningful “olfactory chunks.” Moreover, the parallels with cognitive chunking further underscore an intriguing functional convergence across biological scales. Much like how human memory leverages prior knowledge and statistical regularities to group discrete elements into meaningful clusters, *C. elegans* may similarly exploit inherent statistical structures of odor mixtures to form integrated perceptual representations. This convergence suggests a fundamental neurocomputational principle: when faced with limited processing capacity and complex inputs, biological systems may universally adopt strategies that prioritize holistic, behaviorally relevant representations over detailed elemental fidelity.

## Methods

### Strains

ZAS280*In*[*osm-6*::GCaMP3,*osm-6*::ceNLS-mCherry-2xSV40NLS] ^39–41^.

ZAS325 is a cross between ZAS280 and *unc-31*(e928) ^39^.

ZAS371 is a cross between ZAS280 and *unc-13*(s69) ^39^.

### Worm cultivation

All experiments were performed on 3-days old young adults synchronized by bleaching. Worms were grown at 20°C on NGM plates seeded with OP-50 bacteria.

### Functional imaging and image analysis

Imaging was performed with a Nikon A1R+ confocal laser scanning microscope with a x40 (1.15NA) water immersion objective controlled by the Nikon NIS-elements software. Z-axis intervals were 0.5-0.8μm. Frame rate was ∼1.5 volumes/second. After signal extraction neuronal traces were interpolated to 2 Hz.

Prior to imaging worms were starved for 20 min on un-seeded NGM plates. A single worm was then inserted into a microfluidic chip^42^ and anaesthetized with 10 mM levamisole for an additional 10 min. Imaging began with 30 s of light habituation, followed by 1 min of stimulus presentation, and then 1 min of stimulus removal for a total of 2.5 mins. Stimuli used were diacetyl 10^−4^ (DA), isoamyl alcohol 10^−4^ (IAA), NaCl 50 mM (NaCl), glycerol 1 M (Gly), SDS 0.1% (SDS), quinine 5 mM (Quin), and benzaldehyde 10^−4^ (Benz). In the Ca^2+^ experiments stimuli were not diluted for mixtures. The final concentration of each component in a mixture was the same as of that stimulus alone. The combinations were tested for chemical interactions using SciFinder (https://scifinder-n.cas.org/). No known reactions between components were found. With the exception of the independent IAA+Benz experiment (**Supplementary Figure 9**), neural dynamics responses to individual stimuli and their combinations were measured around the same period of time. The results for the individual stimuli were recently made available ^39^, and here we add the responses to the combinations (**Supplementary Figure 1**).

### Neuron identification and signal extraction

Analysis was done as described previously ^39^. Briefly, neuron tracking was performed with custom software ^73^. Neurons were identified based on anatomical position. All bilateral pairs were pooled aside from the ASE and AWC neurons, the latter of which were classified by activation strength as AWC[strong] and AWC[weak]. Dim, un-segmentable neurons and neurons with strong cross-reads from neighbors were excluded from analysis. Traces were smoothed with a running average of size 5, and step response was calculated as the difference between the average baseline of 10 seconds before the step and the maximum activity in the 7 seconds after the step ^39^. All data analysis was performed with custom MATLAB scripts.

### Fitting models to neuronal integration

Peak neuronal activations were extracted as described in ^39^. Points were constructed as the responses of a neuron to a mixture and its components [X1, X2, y] (**Figure 1B**). Only neurons that responded above a threshold of 0.1 ΔF/F0 to at least one of the stimuli in the triplet were used for the analysis. The principal plane of all points was found using singular value decomposition, and the points were projected onto this plane. First (linear) and second (parabolic) degree polynomials were then fit to the points, weighted by density, and the mean squared error was calculated for both. The points and polynomials were back-projected into the 3D space for plotting. The AWC neurons, which were the only neurons to decrease in response to stimulus presentation, were excluded as outliers (**Figure 1D**).

### Constructing the PCA and the Isomap spaces

Each worm was represented as a 26-dimensional response vector composed of 13 ON step responses and 13 OFF step responses. Bilateral neurons of the same class were pooled except for ASE and AWC.

Due to missing data, and in order to equalize the number of observations for all conditions, worms were randomly resampled with replacement from within each neuron-condition-step combination until 20 observations per condition were obtained^39^. To avoid sampling bias, in all analyses, multiple datasets of 20 worms per condition were independently sampled and the results were aggregated for determining statistical significance.

Before dimensionality reduction, features were z-scored across the resampled dataset. Linear and nonlinear embeddings were obtained for the seven individual stimulus conditions (six odors and control) using PCA and Isomap, respectively. Isomap was fit with Euclidean distances and k = 15 nearest neighbors.

### Statistical and geometric modelling of configurality

For each mixture (A, B, AB) in the 3D PCA space, three Gaussian models were fitted using MATLAB’s fitgmdist function: one to component A (GA), one to component B (GB), and one to the pooled component cloud (A∪B) (G(A∪B)). In all cases, the model used a single Gaussian component with diagonal covariance and regularization value 0.001. Log likelihood was computed for each mixture point x, for each model, to determine how well each cluster explains the mixture. The larger of the two individual component likelihood scores was used for further analysis.

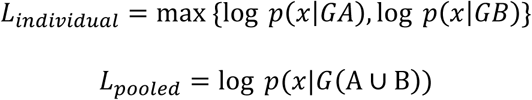

If a mixture lay along the linear axis between its components, it was expected to be better described by the pooled model than by either component alone. The two likelihood distributions were compared across the 20 mixture worms using a one-sided paired Wilcoxon signed-rank test (Tail = left). This procedure was repeated over 100 independent resamplings of 20 worms per condition. p values were adjusted by Benjamini-Hochberg FDR correction.

For the geometric definition of configurality, an ellipse was defined such that the line connecting the two component centroids served as the major axis and the ellipse contour passed through the mixture centroid. Let *A* and *B* denote the component centroids, *M* the mixture centroid, and *C* = 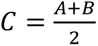 the midpoint between the components. The semi-major axis was defined as *a* = 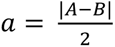.

With the mixture *M* expressed in a 2D coordinate system with x denoting its coordinate along the component axis and y its perpendicular displacement from it, the fitted ellipse satisfied

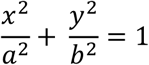

from which the semi-minor axis was obtained as

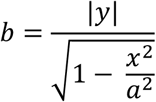

Configurality was then defined as the normalized ellipse aspect ratio

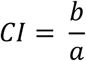

Under this definition, mixtures that lie directly on the component axis yield *CI* = 0, consistent with purely elemental coding, whereas larger off-axis displacements yield larger values of *CI*, indicating stronger configurality.

### Behavioral assays

Choice assays were performed on 3-day old young adult worms synchronized by bleaching and grown on standard NGM plates seeded with OP 50 bacteria. 20 minutes prior to worm loading, 2 µl of 1M NaN_3_ were pipetted onto each of the 4 endpoints on a 9 cm CTX plate (25mM KH2PO4, pH 6.0, 1mM CaCl2,1mM MgSO4, 1.7% agar). Worms were thoroughly washed twice with M9 buffer and once with CTM (5mM potassium phosphate, pH 6.0, 1mM CaCl2,1mM MgSO4), and starved in CTM for 90 minutes.

For IAA and DA, 15 µl of stimulant diluted in ethanol were placed at 2 opposing points and 15 µl of ethanol were placed at the other points. NaCl was applied as described in ^74^. One 5 µl drop of the specified concentration was applied ∼24 hours before the assay, and a second drop ∼6 hours before. To emphasize additivity, and in contrast with the imaging experiments, mixtures in behavioral experiments contained half of the concentrations of the individual stimuli. Mixtures of IAA and DA were made by mixing the odors at a 1:1 ratio. For DA and NaCl mixtures, a solution with half of the NaCl concentration was applied prior to the assay, as specified, and half of the DA concentration was used during the assay.

∼100-200 worms were loaded onto the center of the plate in 15 µl of CTM (which does not contain NaCl), and excess liquid was siphoned off with a rolled up kimwipe. Worms were left to chemotax for 1 hour and then counted. Only fully immobilized worms within 1 cm of the stimulus were counted.

Each experiment included three technical repeats that were averaged for that experiment’s score, and each concentration was tested on at least three different days.

### Analysis of behavioral assays

Chemotaxis index was calculated as the number of worms at both stimulus points minus the number of worms at the control points, divided by the total number of worms on the plate.

Calibration curves were fit using nonlinear least squares (lsqnonlin in MATLAB) as four-parameter hill functions of the form:

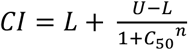

Where L and U are the lower and upper asymptotes, C_50_ is the concentration at half-maximum, and n is the Hill coefficient.

Loewe indices were calculated as dose equivalence ^75^ from the fitted curves as follows:

For a binary mixture of stimuli at concentrations x_mix_ and y_mix_, we measured the behavioral response to the actual mixture as CI_mix_. We then compute the equivalent doses *x_solo_*(*CI_mix_*) and *y_solo_*(*CI_mix_*) that would produce the same effect if each stimulus was presented alone. The Loewe combination index at the observed effect level was then defined as

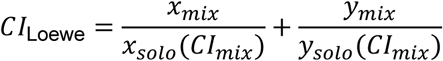

By convention, *CI*_Loewe_ = 1 indicates Loewe additivity (no interaction), *CI*_Loewe_ < 1 indicates synergy, and *CI*_Loewe_ > 1 indicates antagonism.

### Loewe-additive surface (CI = 1)

To visualize the additive expectation for each odor pair, we constructed the Loewe-additive (CI = 1) surface in dose–dose–effect space. For a grid of effect levels *CI* within the range of the two single-odor curves, we computed the single-odor equivalent doses *x_solo_*(*CI_mix_*) and *y_solo_*(*CI_mix_*), and then parameterized the Loewe isobole

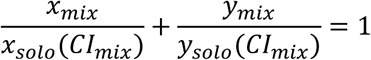

as a line of dose pairs (*x*, *y*) from (0, *y_solo_*(*CI_mix_*)) to (*x_solo_*(*CI_mix_*), 0). Stacking these isoboles over the effect grid yielded a three-dimensional surface of dose pairs and effect levels that are consistent with Loewe additivity, which we refer to as the “Loewe surface”.

### Predicted Loewe effect and distance from the Loewe surface

For each mixture condition (specific doses *x*, *y*), we also computed the effect level predicted by Loewe additivity, *CI*_pred_(*x*, *y*), by numerically solving the isobole equation for *CI_mix_* using the calibrated inverse single-odor curves. The discrepancy between the observed mixture effect and the Loewe prediction was quantified as a vertical distance in effect space,

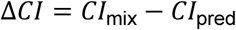

with Δ*CI* > 0 indicating stronger-than-additive responses (synergy) and Δ*CI* < 0 indicating weaker-than-additive responses (antagonism).

## Data availability

All the data, the analyses scripts, as well as the scripts that produce the figures are available at: https://github.com/zaslab/Nonlinear-sensory-dynamics-drive-configural-percepts Complete raw data figures are provided in Supplementary Figures 10-40.

