## Supplementary File for "Nonlinear sensory integration drives configural olfactory representations"

### Supplementary Notes, Table, and Figures

#### Supplementary Note 1. Linear and nonlinear structures encode different features of the environment.

The principal components of the manifold are believed to correspond to perceptual properties of stimuli (*e.g.,* the first component often reflects pleasantness, though the second varies across studies [8,9]). Therefore, the location of a stimulus cluster in the perceptual space should, in principle, correlate with features of the stimulus that have some relevance to the organism. We applied an SVM classifier to analyze the linear separability of the stimuli in both the PC space and within a nonlinear manifold. For each perceptual space we attempted to separate the stimuli by their identity and by their valence, with IAA, DA, and NaCl being grouped as attractive, and SDS, quinine and glycerol as aversive.

Surprisingly, while the identities of the stimuli are better encoded in the linear PCA space, the valence was best separated on the nonlinear manifold (**Supplementary Figure 4C-G**). Separability by identity saturated at 5-6 dimensions, with the linear space providing consistently better accuracy than the nonlinear manifold. Including additional dimensions decreased accuracy, likely due to overfitting. Interestingly, separation by valence required fewer dimensions to reach saturation in the nonlinear than in the linear space. This suggests that the representations of stimulus identities in the sensory neurons are high-dimensional and linear, while the more abstract feature of valence is encoded in a low-dimensional, nonlinear manifold.

The observation that in this perceptual space mixtures tend to fall between the component clusters, regardless of the spread of the components, suggests a degree of convergence of the percepts. “Olfactory white”, a phenomenon in which as mixtures increase in number of components they tend to converge on a similar and neutral perceptual experience has been previously described in humans [10]. To test if a similar phenomenon exists in the neuronal perceptual space of *C. elegans*, we calculated the Euclidean distances between all clusters including 7 single components, 6 binary mixtures, and 2 triple mixtures. We found that the pairwise distances decrease as the number of components increases, converging onto the center of the perceptual space where the neutral control cluster lies (**Supplementary Figure 4H**).

### Supplementary Note 2. Contexts that produce irrational decision-making are enriched for configural mixtures

*C. elegans* emerges as a valuable model for studying principles governing various decision-making behaviors, including rationality and effort discounting [1,11–14]. Here, we studied whether the neurally-defined measure of configurality corresponds to rational decision-making. Following rational choice theory, we posited that each stimulus holds an absolute value for the worm. Elementally perceived mixtures, where components are perceived independently, should preserve these absolute values, leading to consistent, transitive preferences. In contrast, configurally perceived mixtures, which generate a novel percept, would create a new, context-dependent value, thereby disrupting the stable hierarchy and enabling intransitive, or "irrational", choices.

We tested this by re-analyzing the comprehensive behavioral dataset from [1], which scored the rationality of the preference hierarchy of thousands of stimulus triplets based on their adherence to transitivity. We aggregated all triplets from this dataset that contained the components of our identified mixtures: the highly configural DA+IAA and the elemental IAA+NaCl and DA+NaCl, and compared the probability distributions of these groups. We found that triplets containing the configural DA+IAA pair were significantly less likely to be transitive than triplets containing the elemental mixtures (**Supplementary Figure 8A**). Breaking down the triplets by the severity of the violation of transitivity showed that the DA+IAA pair is significantly enriched among triplets with high probability of violating moderate stochastic transitivity (**Supplementary Figure 8B**). These results suggest that when required to choose between options that form configural percepts when presented together, violations of rational decision-making are more likely.

| | In-sample RMSE | Mean CV RMSE | Std CV RMSE | BIC | Parameters |
|---|---|---|---|---|---|
| Linear | 0.133 | 0.137 | 0.028 | -318.22 | 2 |
| Quadratic | 0.106 | 0.109 | 0.019 | -351.17 | 3 |
| Cubic | 0.105 | 0.11 | 0.019 | -348.29 | 4 |

**Supplementary Table 1. Metric comparison for linear, quadratic, and cubic model fits.**

The quadratic model explains the data best with the lowest BIC. Although a cubic model performed similarly to the quadratic, it was disfavored by BIC, indicating that the quadratic model provided the best balance between accuracy and complexity.

### Supplementary Figures

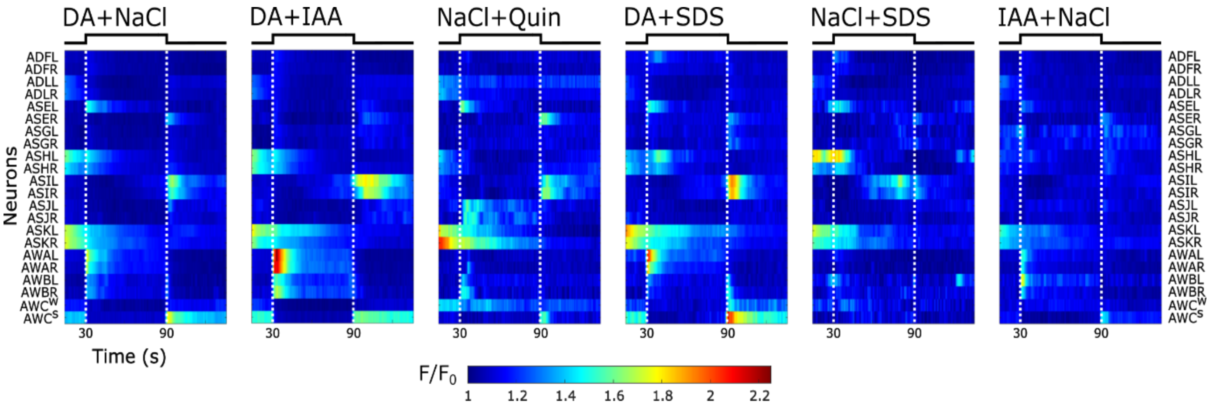


**Supplementary Figure 1. Heat maps of the 22 amphid neuron responses to binary mixtures.**

The mixtures were added at t=30 sec (ON step) and removed at t=90 sec (OFF step) as indicated by the white dashed lines. Color bar indicates neural activity. The bilateral AWC neurons are depicted as AWC$^{S}$ (strong) and AWC$^{W}$ (weak), depending on their response magnitude in each animal, rather than R (right) or L (left) as shown for the other neurons.

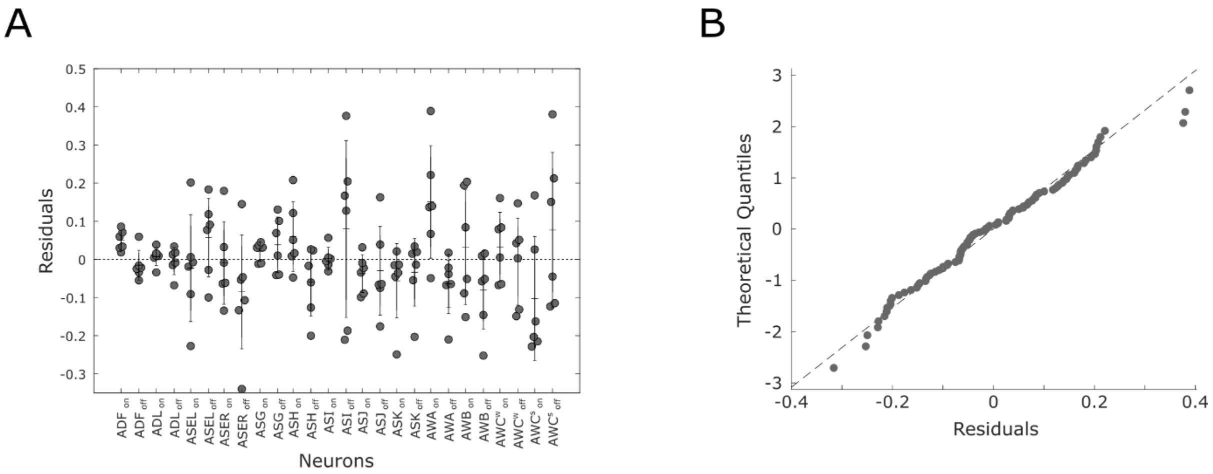


**Supplementary Figure 2.**

**(A)** Residual plot of the linear model by neuron for both on and off steps (indicated next to the neuron name). Bars represent standard deviation.

**(B)** Q-Q plot showing that the residuals are normally distributed.

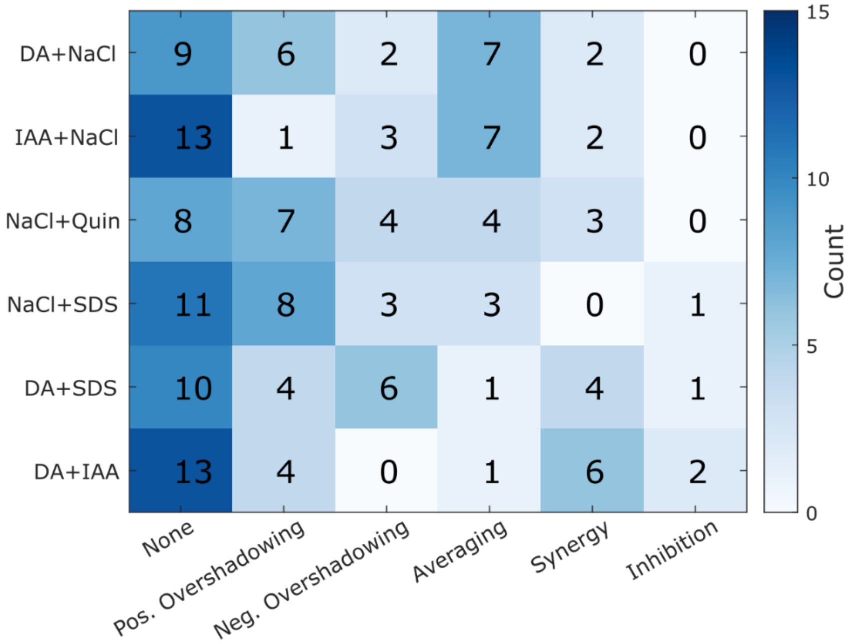


**Supplementary Figure 3.** Occurrence of integration types by mixture. None - Neuron did not respond to either component or to the mixture. Pos. overshadowing – mixture response is similar to the stronger component ($A > B, AB = A$). Neg. overshadowing – mixture response is similar to the weaker component ($A > B, AB = B$). Averaging – mixture response is between both components ($A > AB > B$). Synergy – Mixture response is stronger than either component ($AB > \{A, B\}$). Inhibition – mixture response is weaker than either component ($AB < \{A, B\}$). For any mixture, neurons that did not respond to either component or the mixture with an amplitude above 0.1 ΔF/F0 were excluded. Only neurons with significant differences were included (two-tailed t test). For example, a response was considered synergistic only if both $AB > A$ and $AB > B$ were significant.

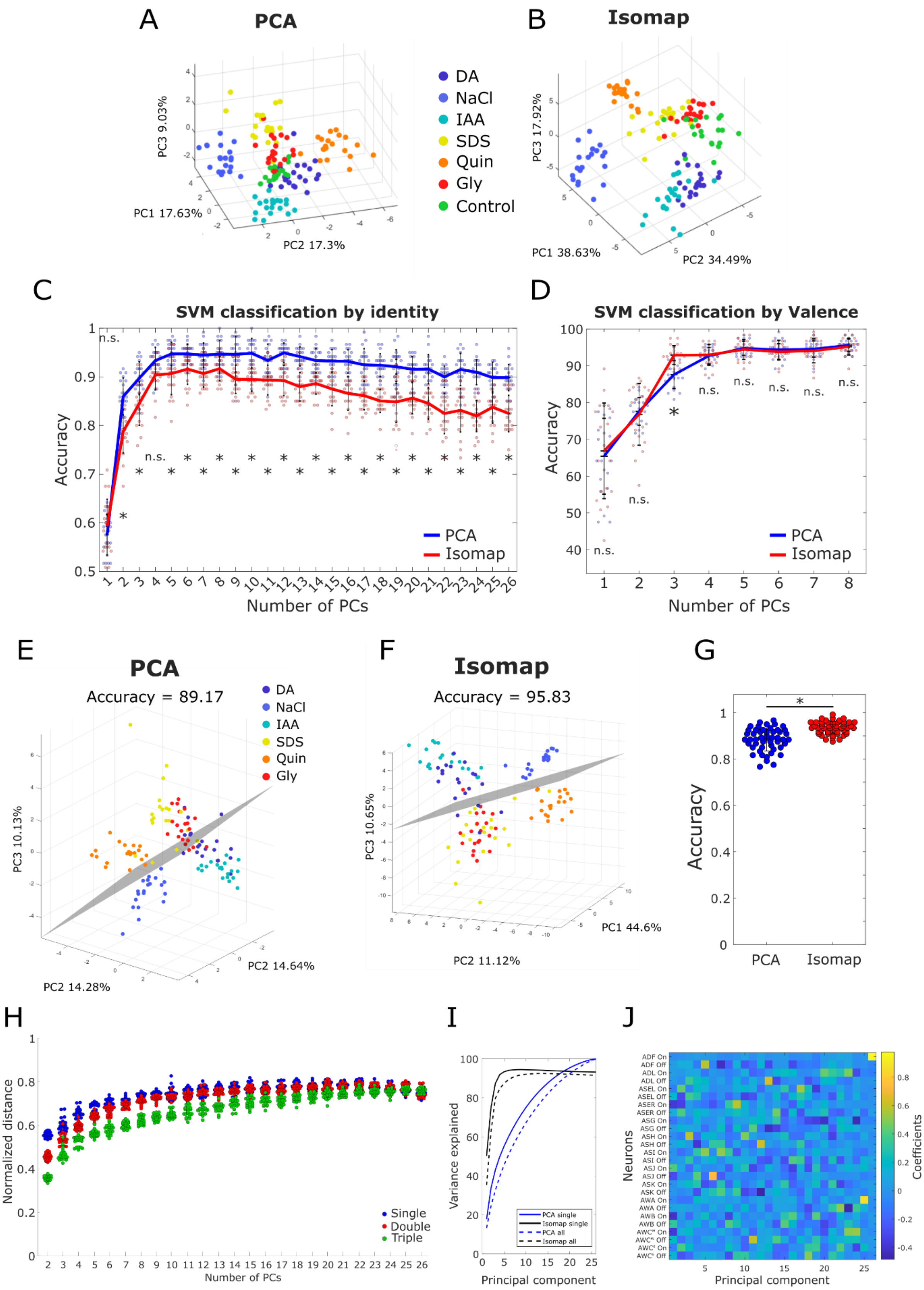
A
PCA
PC3 9.03%
PC1 17.63%
PC2 17.3%
DA
NaCl
IAA
SDS
Quin
Gly
Control
B
Isomap
PC3 17.92%
PC1 38.63%
PC2 34.49%
C
SVM classification by identity
Accuracy
Number of PCs
n.s.
PCA
Isomap
D
SVM classification by Valence
Accuracy
Number of PCs
n.s.
PCA
Isomap
E
PCA
Accuracy = 89.17
PC3 10.13%
PC2 14.28%
PC2 14.64%
DA
NaCl
IAA
SDS
Quin
Gly
F
Isomap
Accuracy = 95.83
PC3 10.65%
PC1 44.6%
PC2 11.12%
G
Accuracy
PCA
Isomap
H
Normalized distance
Number of PCs
Single
Double
Triple
I
Variance explained
Principal component
PCA single
Isomap single
PCA all
Isomap all
J
Neurons
Principal component
Coefficients

**Supplementary Figure 4. Identity and valence are differently encoded in the neural space.**

**(A-B)** Scatter plots of system responses to individual stimuli in the first three components of PCA (replicated from **Figure 3A** and shown here for convenience) **(A)** and Isomap **(B)** spaces. Each point is a re-sampled worm (see methods).

**(C)** Accuracy scores of linear SVM classifiers trained on stimulus identities using different numbers of principal components. Lines represent the means of 20 random resamplings of worms. Accuracy scores were obtained from 3-fold cross-validation.

**(D)** Accuracy scores of linear SVM classifiers trained on valence using different numbers of principal components. DA, IAA and NaCl were marked as attractive, Gly, SDS and Quin were marked as aversive. Lines represent the means of 20 random resamplings of worms. Accuracy scores were obtained from 3-fold cross-validation.

**(E-G)** Representative plots of the SVM classification by valence with 3 components of PCA **(E)** and Isomap **(F)**.

**(H)** Mixture percepts converge as the number of components increases. Euclidean distances between mixtures with different numbers of components. Each point represents the mean pairwise distances of all clusters in a random resampling of 20 worms per stimulus, grouped by mixture complexity - blue: single stimulus, red: binary mixtures, green, triple mixture. Distances were normalized by the largest pairwise distance in the sample.

**(I)** Variance explained by the principal components of PCA and Isomap when performed on only single stimuli (solid lines), and including the mixtures (dashed lines)

**(J)** PC loadings. Contributions of each neuron’s ON and OFF step responses to principal components.

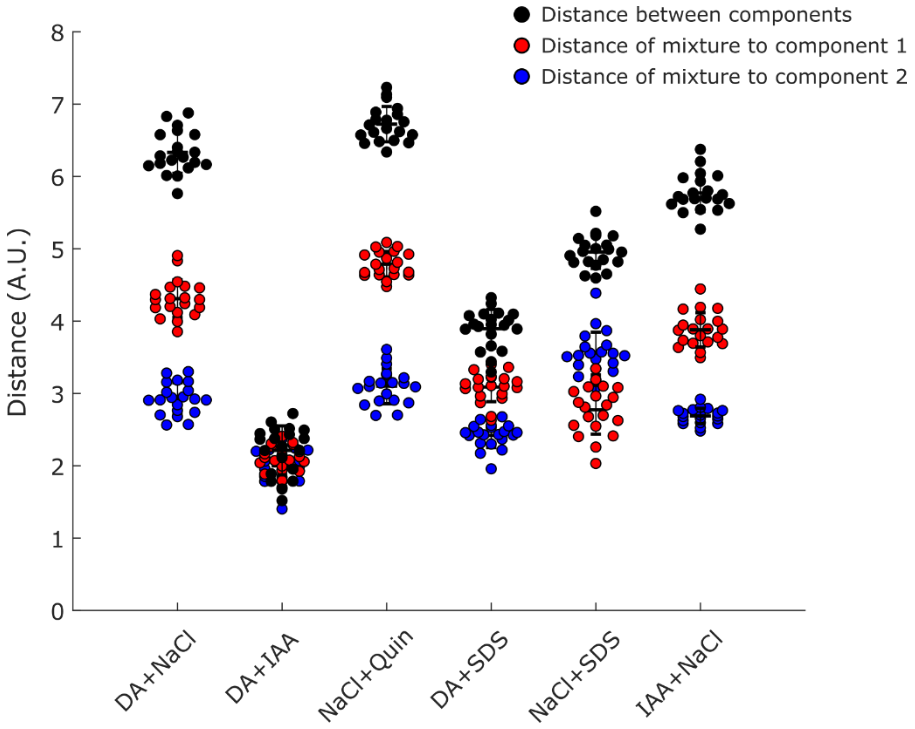


**Supplementary Figure 5. Mixtures in PC space are found between the component clusters.** In all tested mixtures the distance between the two components (black) is greater than the distances between each of the components and the mixture (red/blue).

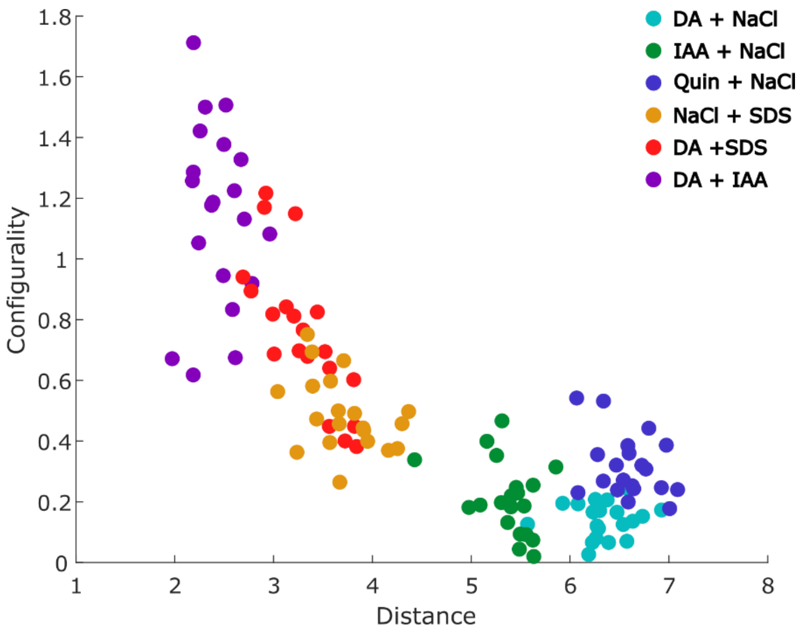


**Supplementary Figure 6.** The configurality metric plotted against the Euclidean distance between components in the neural space. Mixtures, for which the individual components are more similar to one another (lower distance on the x axis) show greater configurality. For example, the coding patterns of IAA and DA, two attractive odorants, are very close in the PC space, yet their combination results in the highest configural coding.

A

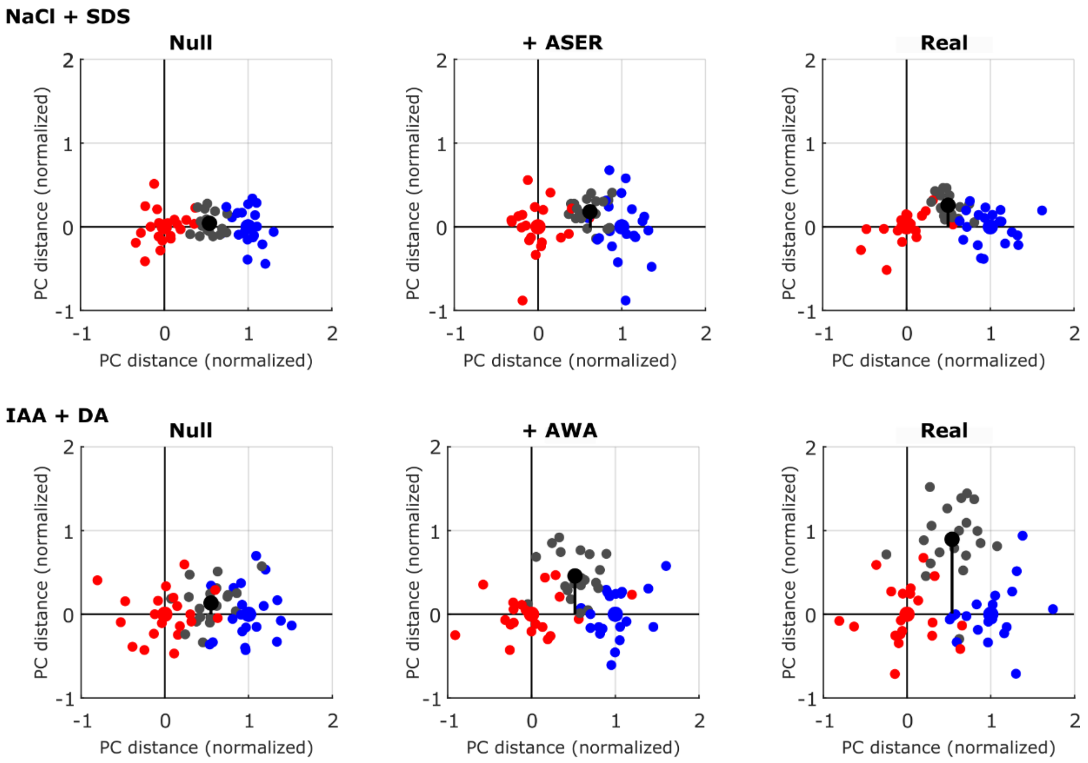


B

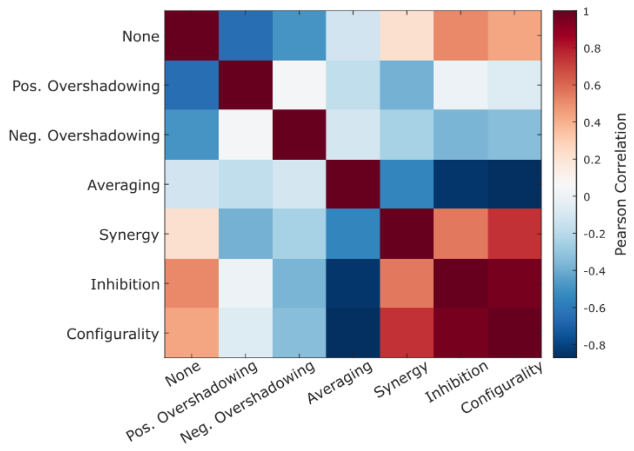


**Supplementary Figure 7. The nonlinear synergy and inhibition integration types are strongly correlated with configurality.**

**(A)** 2D projections of Figure 4D-E. Examples of contribution to configurality by a single neuron integrating by inhibition (top) and synergy (bottom). Component clusters are in red and blue, mixture cluster in black. Distances were scaled to make the centerline be of length 1. Left: Null mixture (mean of components), middle: null mixture with a single real measured neuron, right: full measured mixture.

**(B)** Correlations between the number of occurrences of integration types and the configurality score.

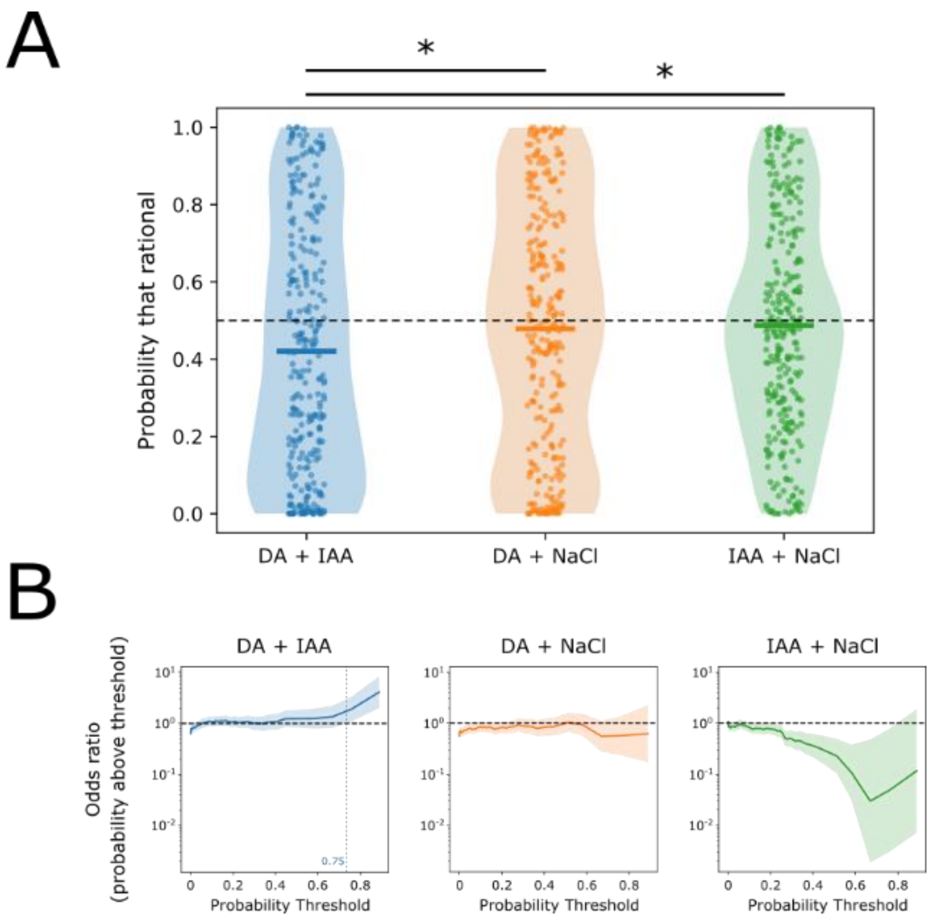


**Supplementary Figure 8. Choices that form a configural percept may violate the transitivity paradigm and lead to irrational decision-making.**

**(A)** Mean and distribution of "Probability that rational" for all stimulus triplets that contain the configural DA+IAA mixture vs the elemental DA+NaCl, and IAA+NaCl mixtures. Dashed line marks probability of 0.5. Data was obtained from [1]. Asterisks indicate significant differences in both the mean (Welch's t-test: Blue vs Orange, p = 0.032, Blue vs Green, p = 0.009) and the shape of the distribution (Two-sample Kolmogorov-Smirnov test: Blue vs Orange, p = 0.013, Blue vs Green, p = 0.003).

**(B)** Odds ratios as a function of threshold for "Probability that violates moderate stochastic transitivity" for triplets that contain the indicated pair versus all other triplets. Values above the horizontal dashed line indicate enrichment. Shaded region marks the 95% confidence interval. The vertical dashed line in DA+IAA indicates the probability above which there is sustained significant enrichment of DA+IAA.

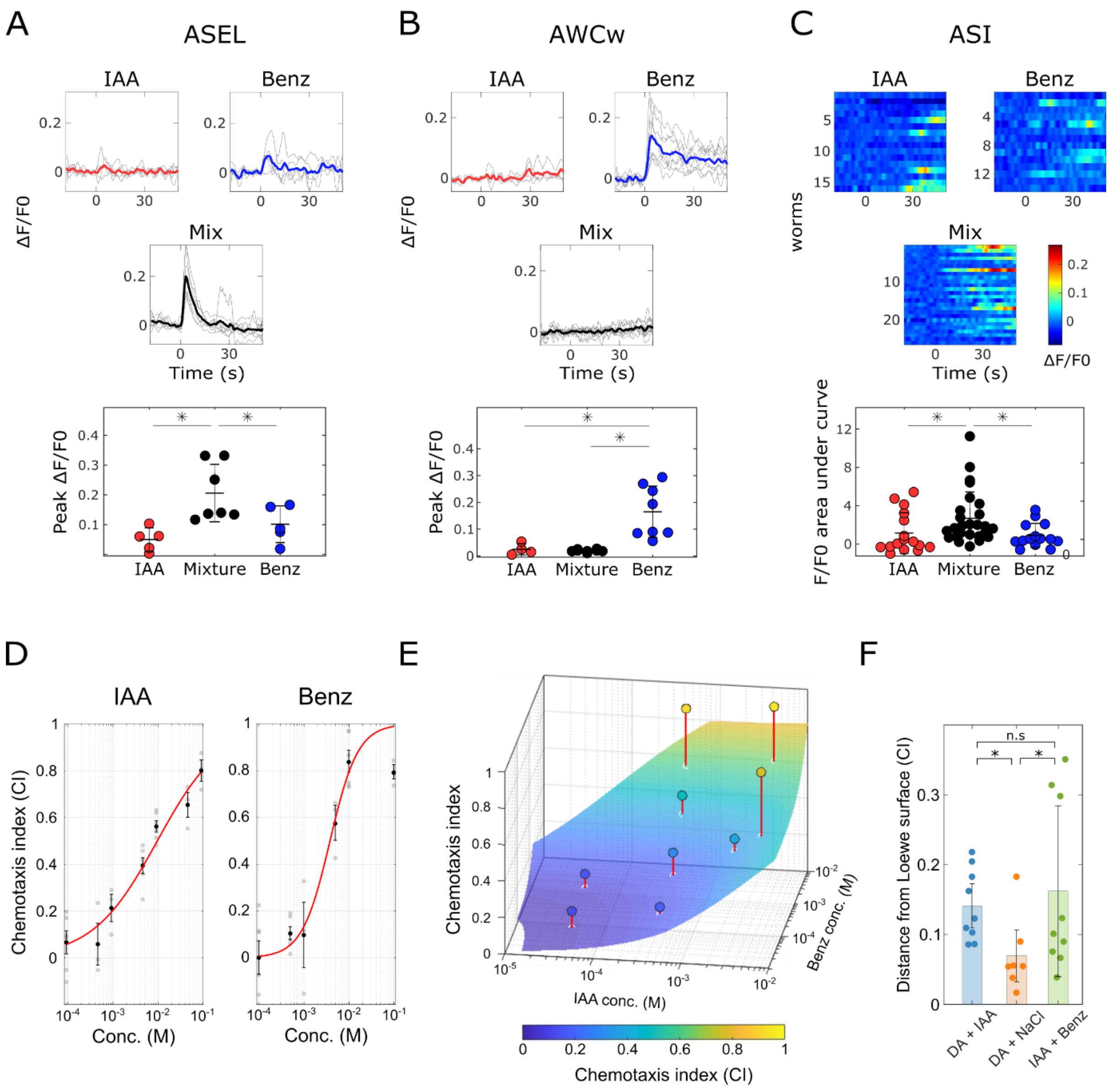


**Supplementary Figure 9. A mixture of the attractive food odors IAA and Benz invokes strong nonlinear neuronal integration and behavioral synergy**

**(A-C)** Synergy and inhibition in the integration of the IAA+Benz mixture in individual neurons. Emergent activity in ASEL **(A)**, inhibitory effect in AWC$_w$ **(B)**, synergy in ASI **(C)**. Top: Single neuron traces (grey lines in A, B and heatmap rows in C) and population mean traces of the responses. Red and blue: individual stimulus components; black: their mixture. Bottom: Peak response amplitudes in A, B and area under the curve in C (ΔF/F0). * $p<0.05$.

**(D)** Dose-response calibration curves for the individual components. Black points represent averages of technical repeats. Black bars are standard error of the mean for all repeats of the same concentration. Red curves represent the fitted logistic model.

**(E)** Predicted chemotaxis under Loewe dose additivity based on the calibration curves in (D). The colored surface represents the expected Loewe response $CI_L$ for each pair of doses, computed from the isobole equation $\frac{x}{c_A^{\text{eq}}(CI_{mix})} + \frac{y}{c_B^{\text{eq}}(CI_{mix})} = 1$ (see methods). Filled circles mark the measured mixture responses; red stems indicate the vertical deviation from the additive prediction surface (positive = synergy, negative = antagonism).

**(F)** Summary of deviations from additivity for IAA+Benz. IAA+DA and DA+NaCl bars are reproduced from **Figure 5**. DA+IAA vs IAA+Benz p = 0.264. DA+NaCl vs IAA+Benz p = 0.026

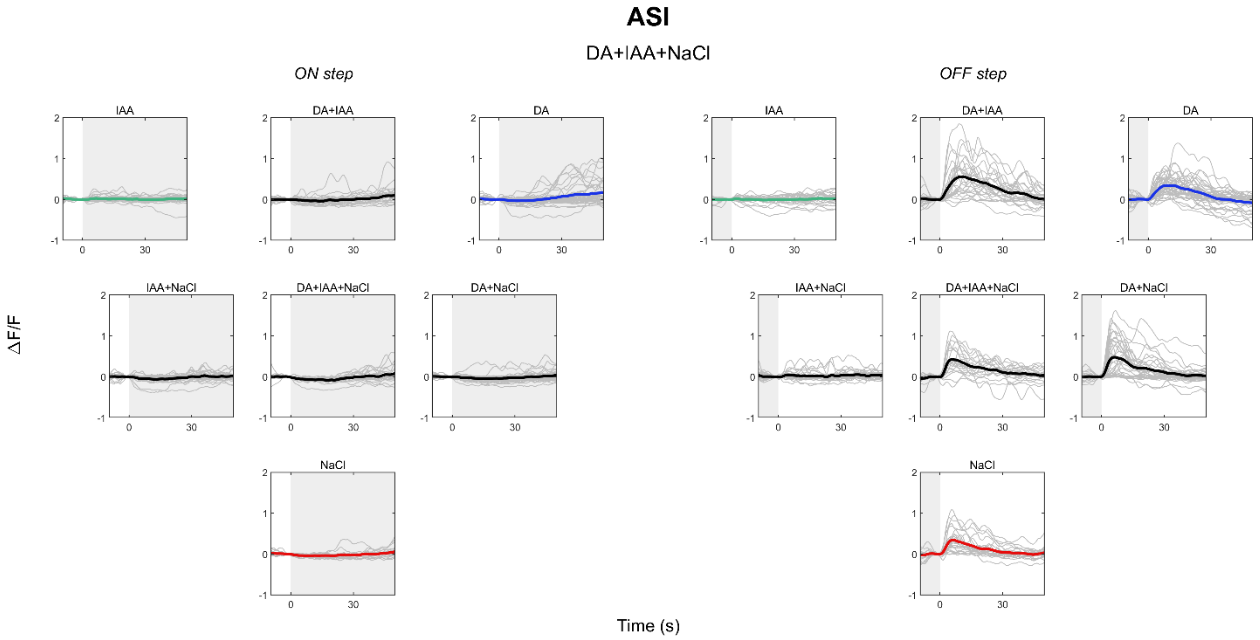


**Supplementary Figure 10.** Raw mean traces of ASI in DA (blue), IAA (green) and NaCl (red), and their mixtures (black). Individual neuron traces in grey. Shaded area indicates presence of stimulus.

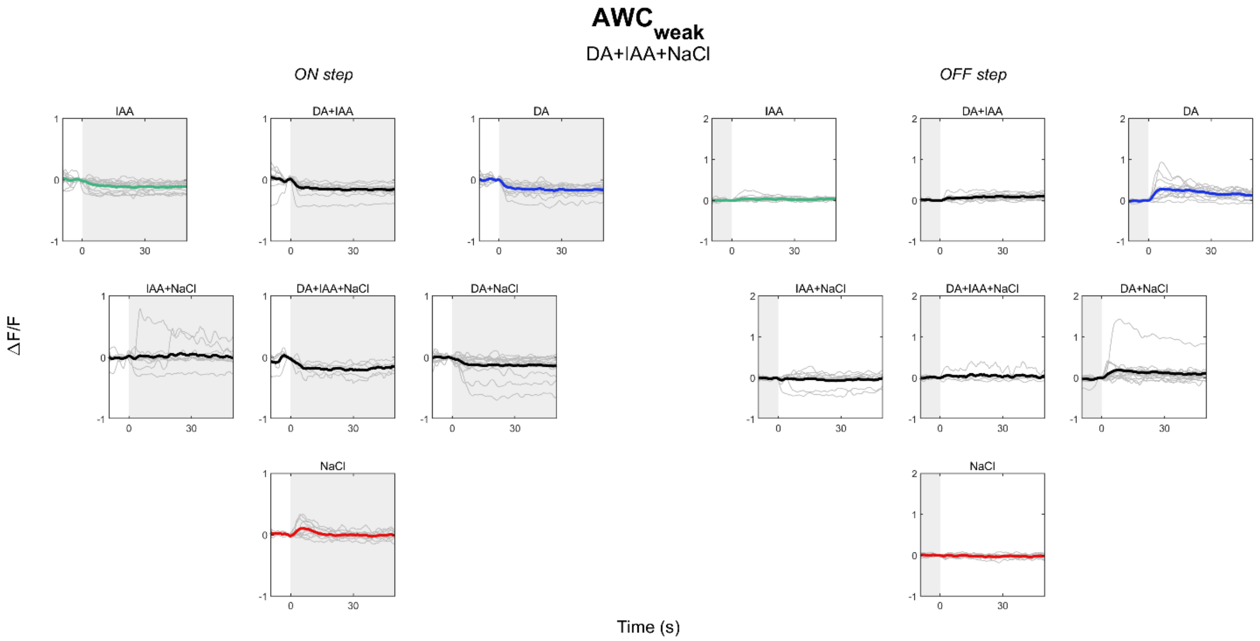


**Supplementary Figure 11.** Raw mean traces of $AWC_{weak}$ in DA (blue), IAA (green) and NaCl (red), and their mixtures (black). Individual neuron traces in grey. Shaded area indicates presence of stimulus.

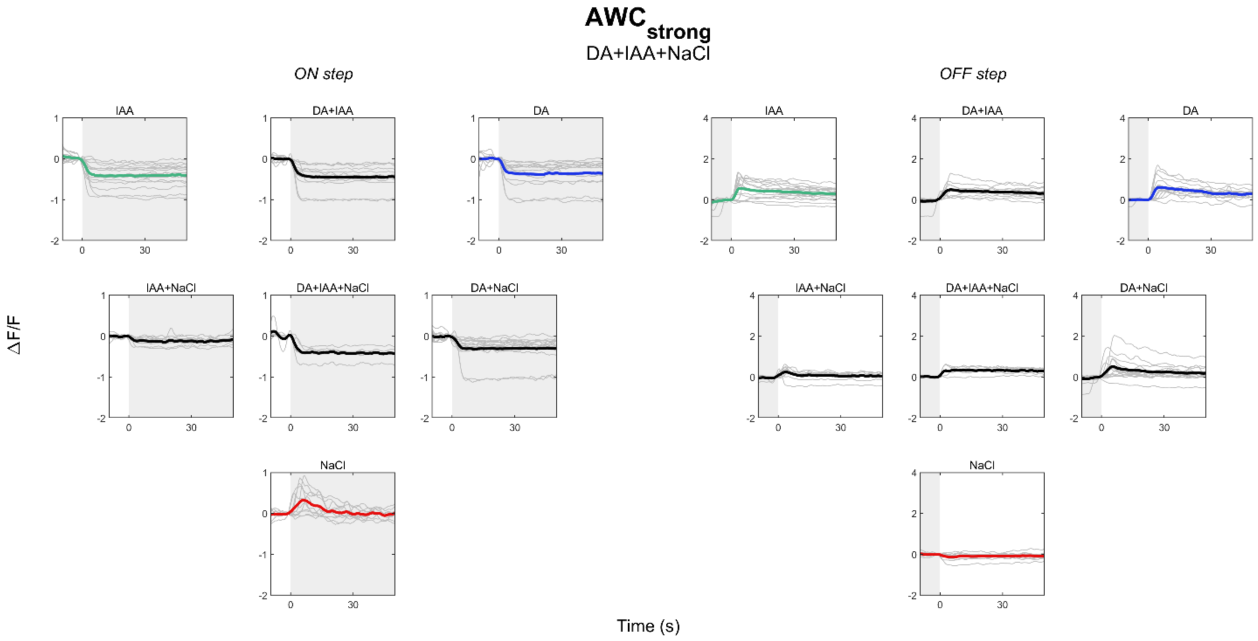


**Supplementary Figure 12.** Raw mean traces of AWC$_{strong}$ in DA (blue), IAA (green) and NaCl (red), and their mixtures (black). Individual neuron traces in grey. Shaded area indicates presence of stimulus.

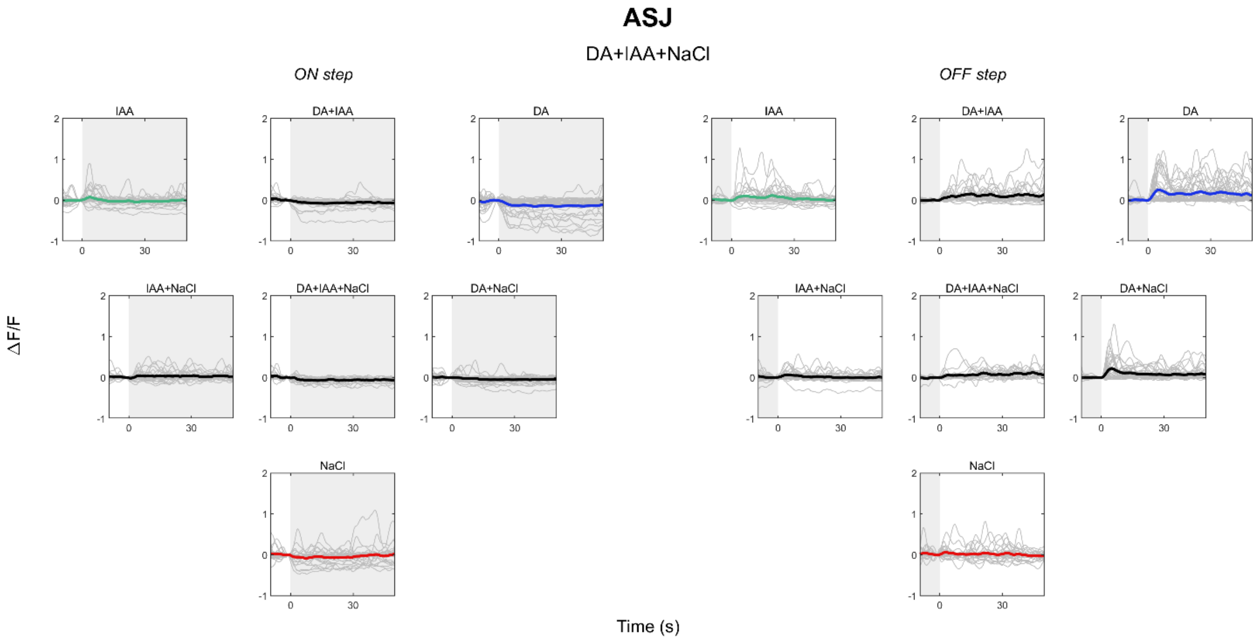


**Supplementary Figure 13.** Raw mean traces of ASJ in DA (blue), IAA (green) and NaCl (red), and their mixtures (black). Individual neuron traces in grey. Shaded area indicates presence of stimulus.

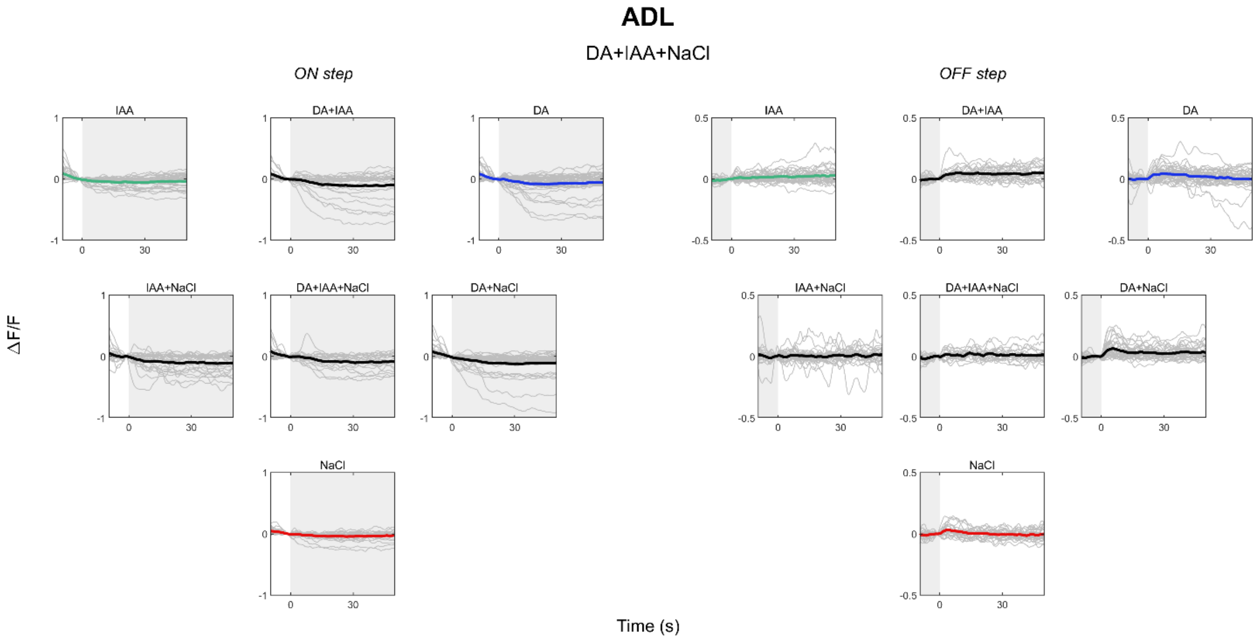


**Supplementary Figure 14.** Raw mean traces of ADL in DA (blue), IAA (green) and NaCl (red), and their mixtures (black). Individual neuron traces in grey. Shaded area indicates presence of stimulus.

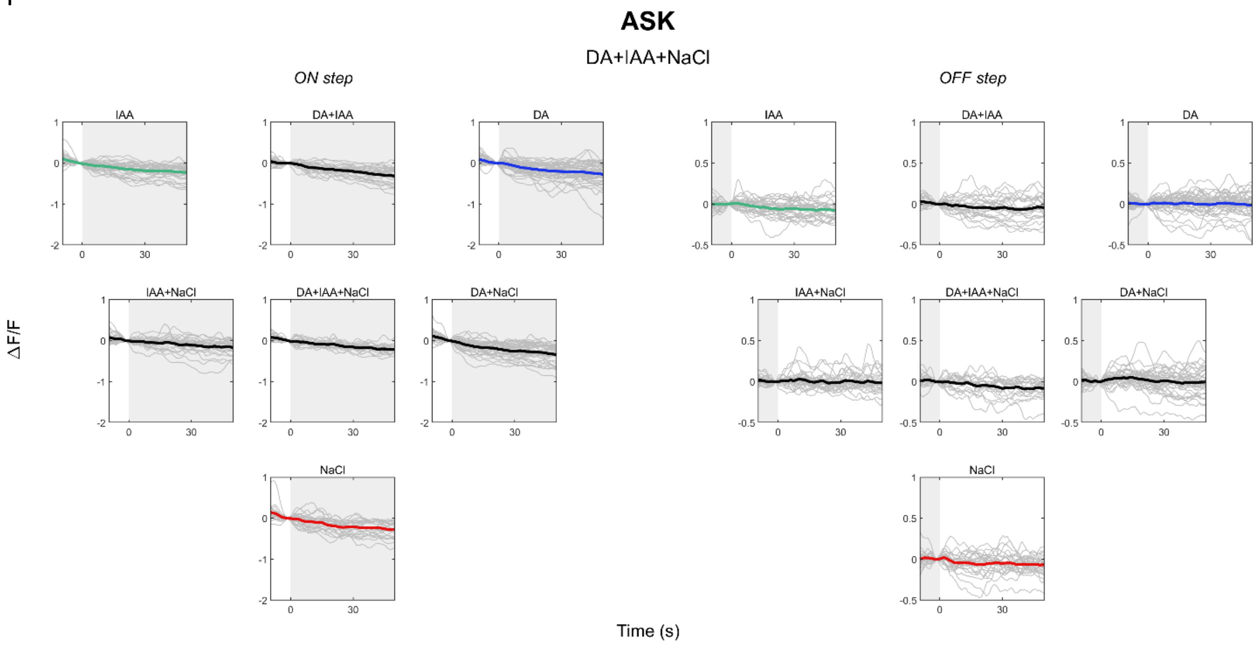


**Supplementary Figure 15.** Raw mean traces of ASK in DA (blue), IAA (green) and NaCl (red), and their mixtures (black). Individual neuron traces in grey. Shaded area indicates presence of stimulus.

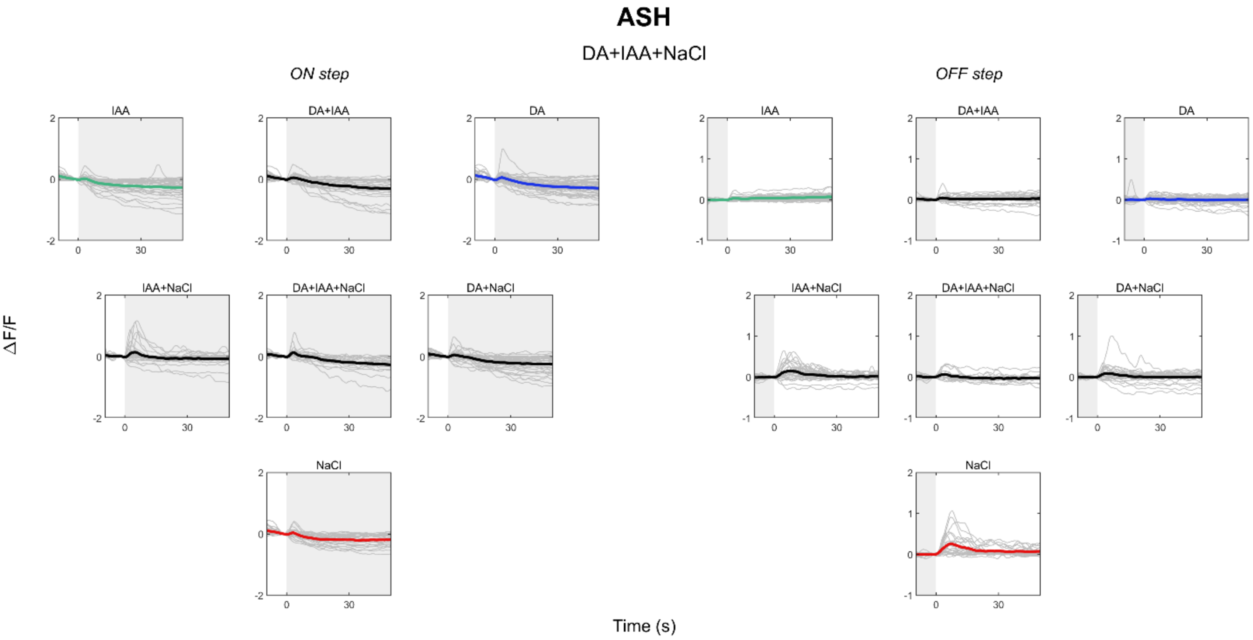


**Supplementary Figure 16.** Raw mean traces of ASH in DA (blue), IAA (green) and NaCl (red), and their mixtures (black). Individual neuron traces in grey. Shaded area indicates presence of stimulus.

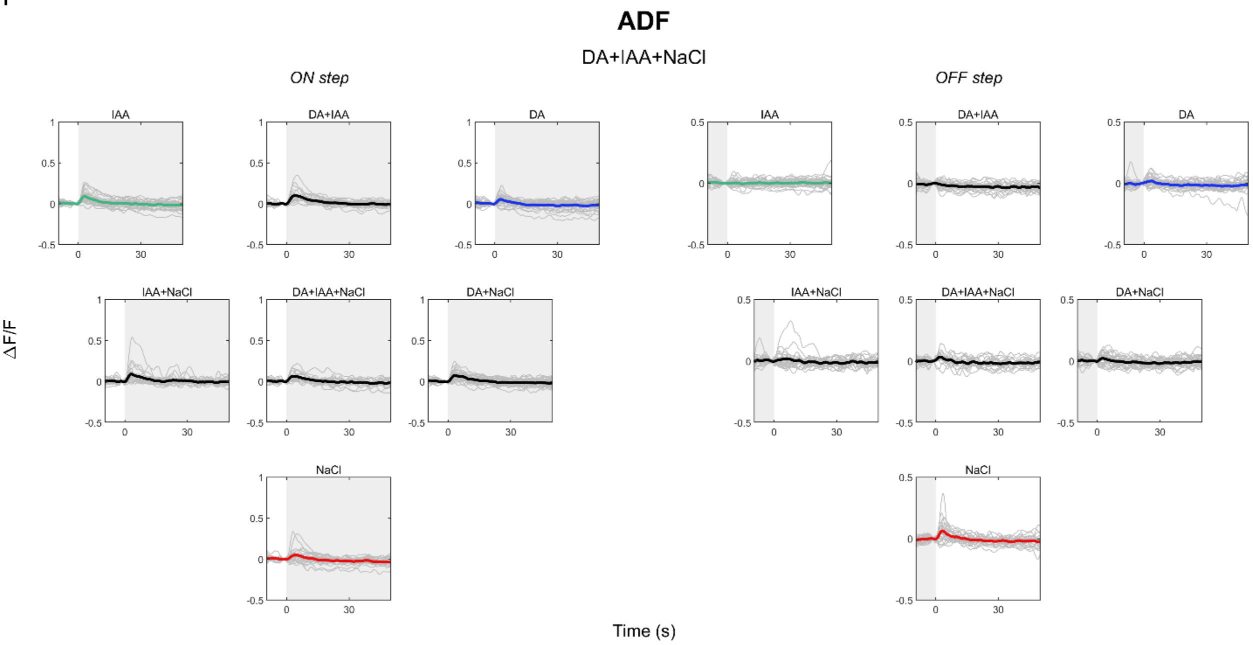


**Supplementary Figure 17.** Raw mean traces of ADF in DA (blue), IAA (green) and NaCl (red), and their mixtures (black). Individual neuron traces in grey. Shaded area indicates presence of stimulus.

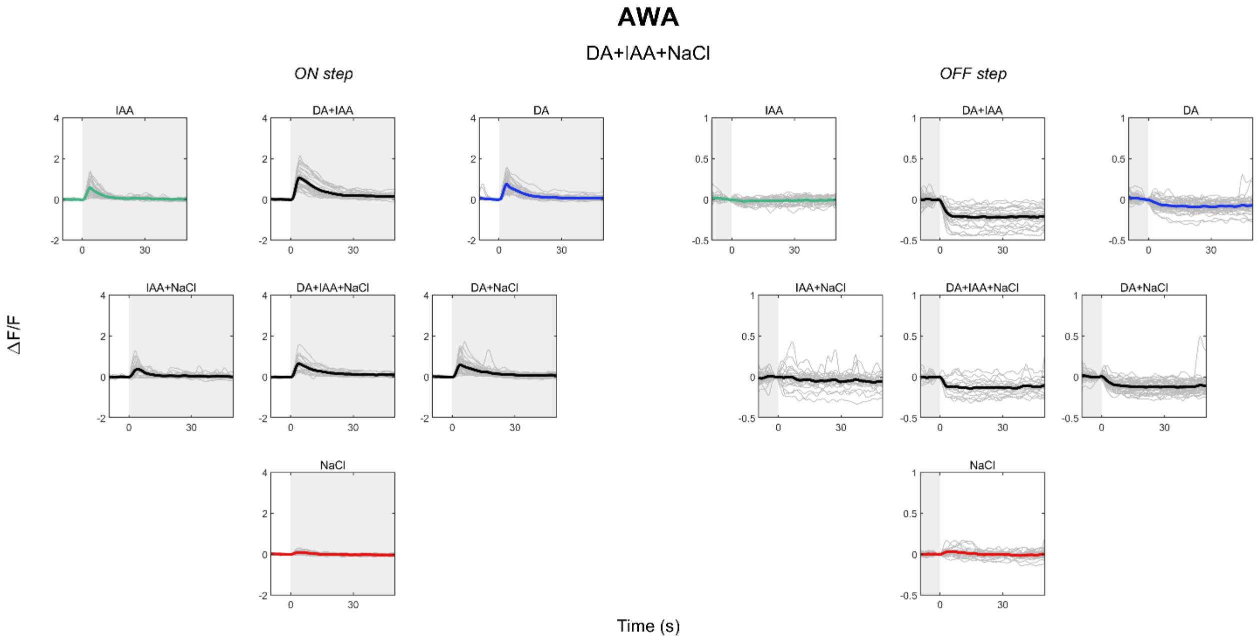


**Supplementary Figure 18.** Raw mean traces of AWA in DA (blue), IAA (green) and NaCl (red), and their mixtures (black). Individual neuron traces in grey. Shaded area indicates presence of stimulus.

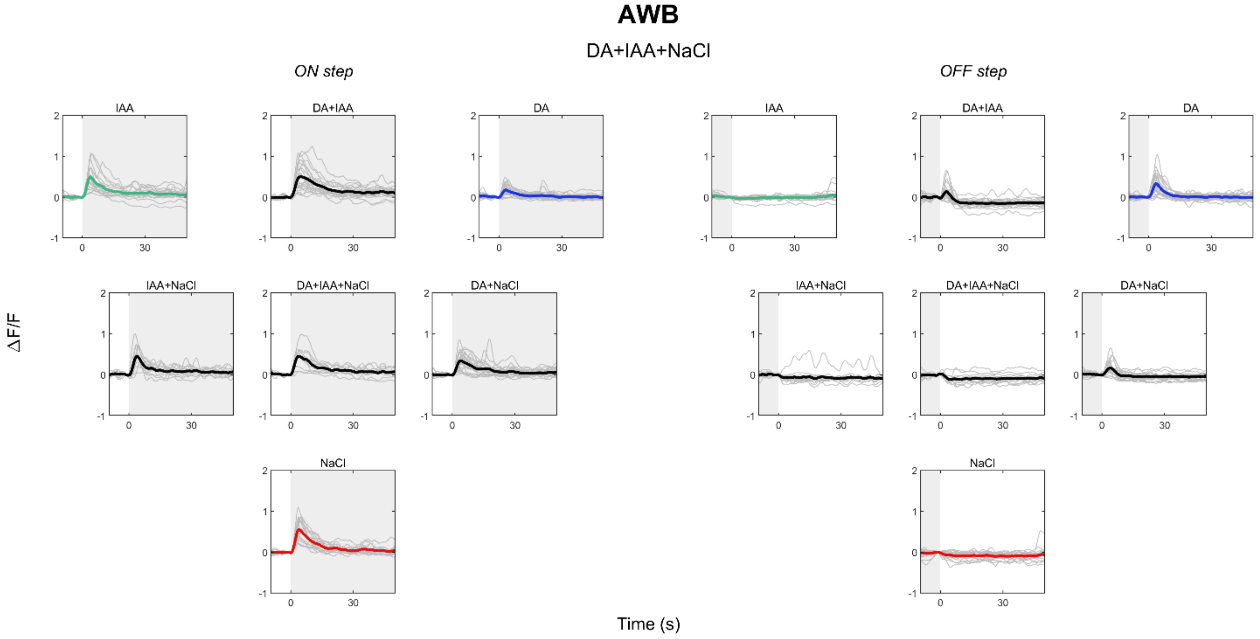


**Supplementary Figure 19.** Raw mean traces of AWB in DA (blue), IAA (green) and NaCl (red), and their mixtures (black). Individual neuron traces in grey. Shaded area indicates presence of stimulus.

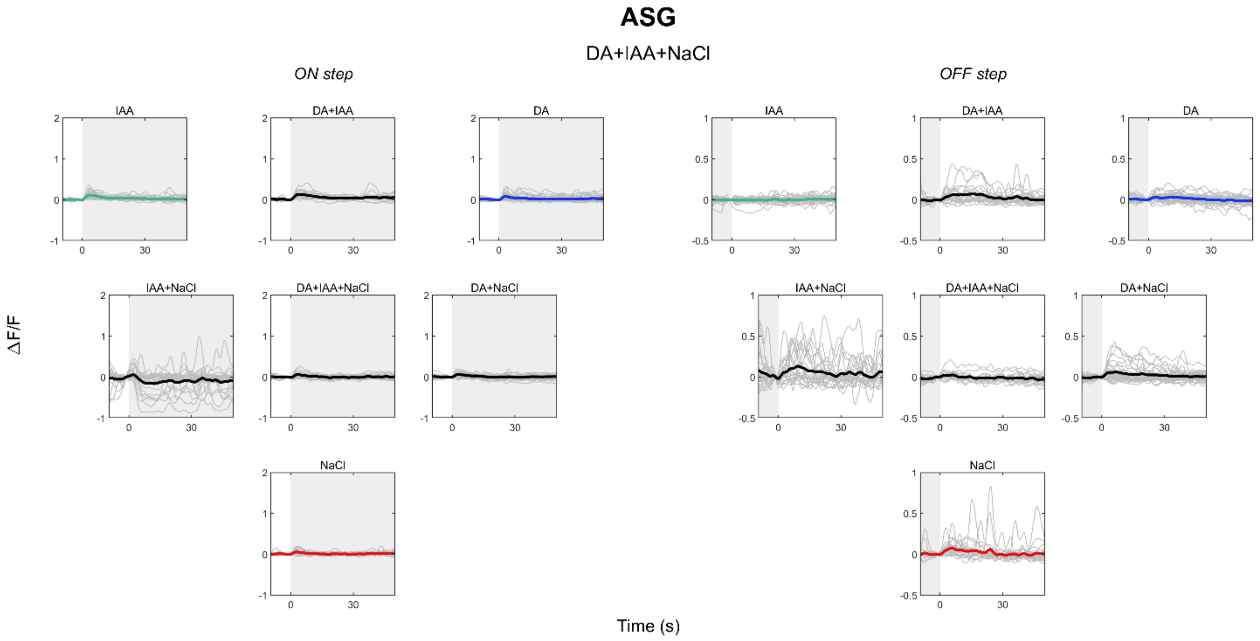


**Supplementary Figure 20.** Raw mean traces of ASG in DA (blue), IAA (green) and NaCl (red), and their mixtures (black). Individual neuron traces in grey. Shaded area indicates presence of stimulus.

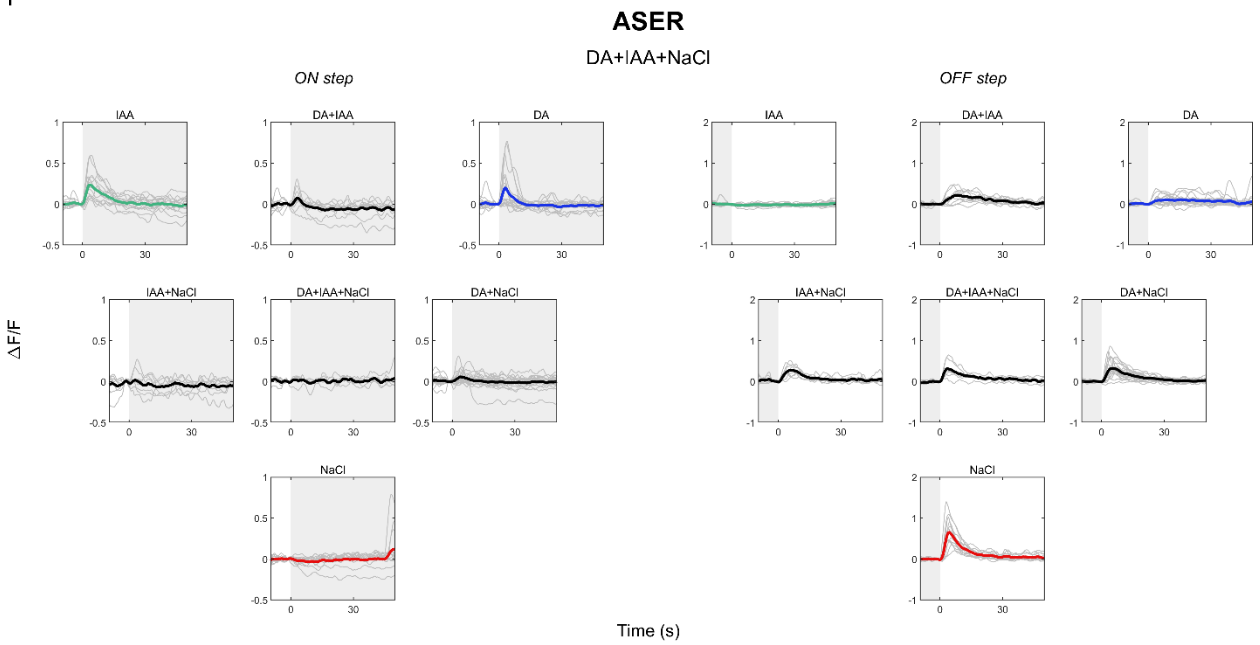


**Supplementary Figure 21.** Raw mean traces of ASER in DA (blue), IAA (green) and NaCl (red), and their mixtures (black). Individual neuron traces in grey. Shaded area indicates presence of stimulus.

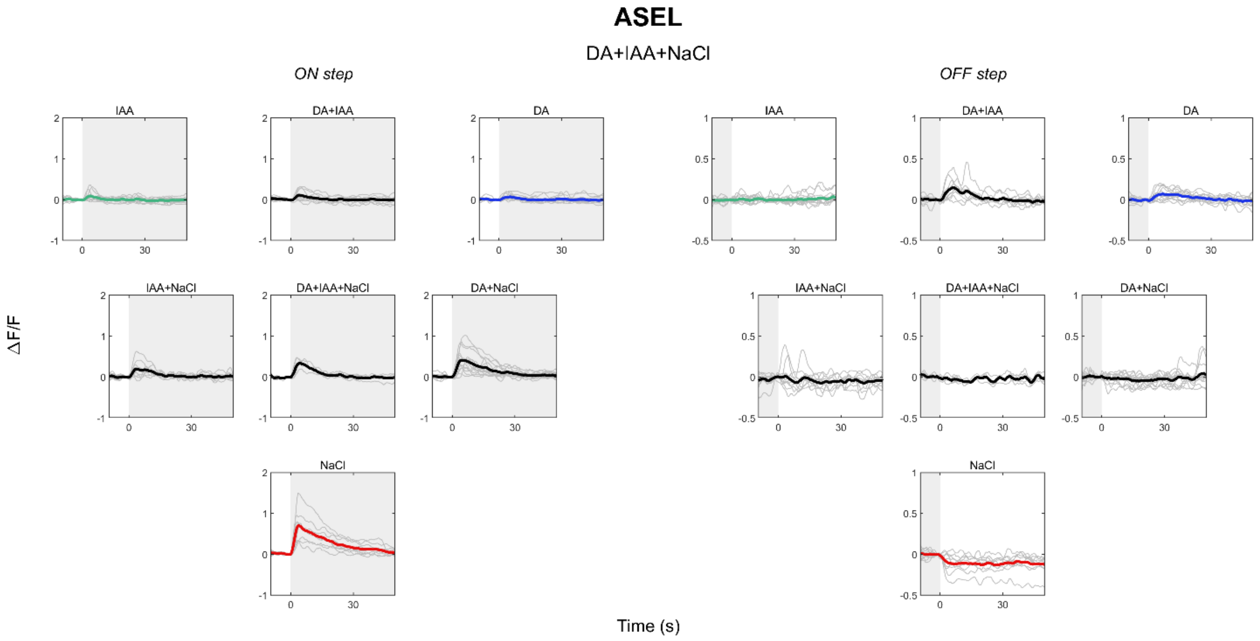


**Supplementary Figure 22.** Raw mean traces of ASEL in DA (blue), IAA (green) and NaCl (red), and their mixtures (black). Individual neuron traces in grey. Shaded area indicates presence of stimulus.

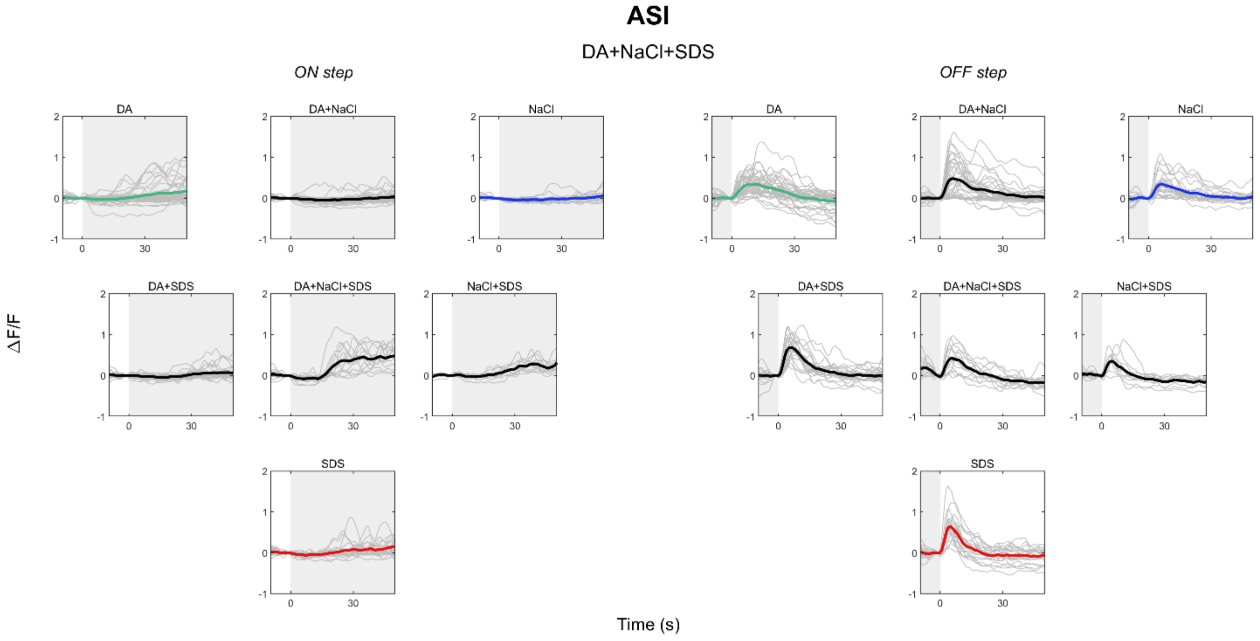


**Supplementary Figure 23.** Raw mean traces of ASI in NaCl (blue), DA (green) and SDS (red), and their mixtures (black). Individual neuron traces in grey. Shaded area indicates presence of stimulus.

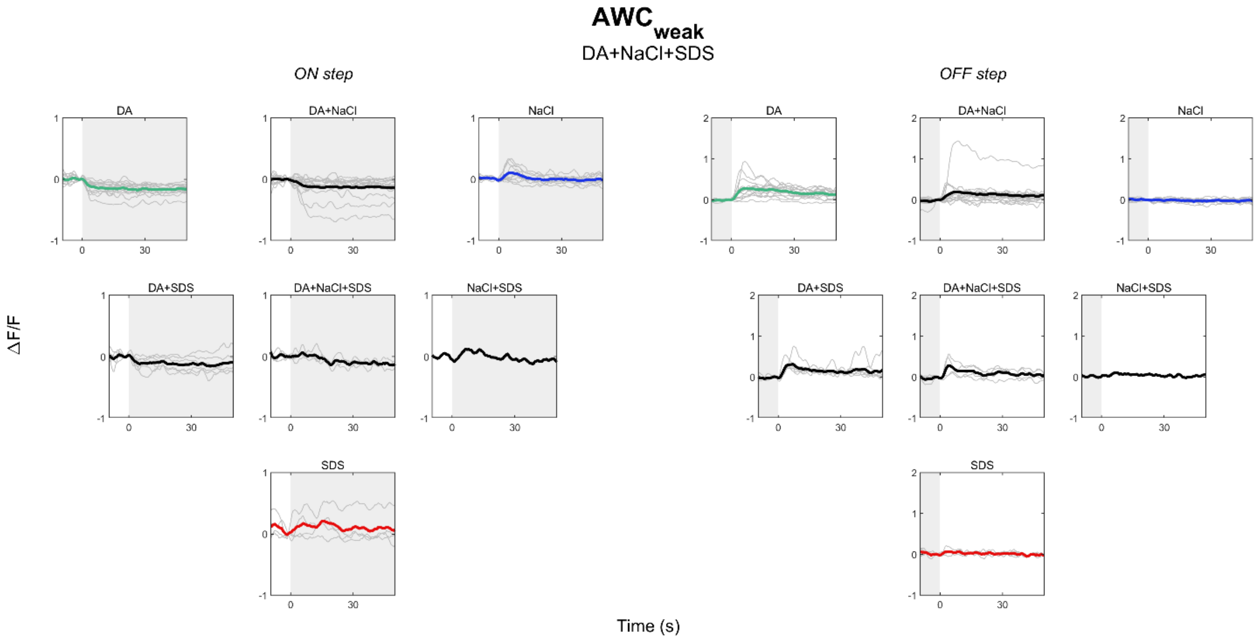


**Supplementary Figure 24.** Raw mean traces of AWC$_{weak}$ in NaCl (blue), DA (green) and SDS (red), and their mixtures (black). Individual neuron traces in grey. Shaded area indicates presence of stimulus.

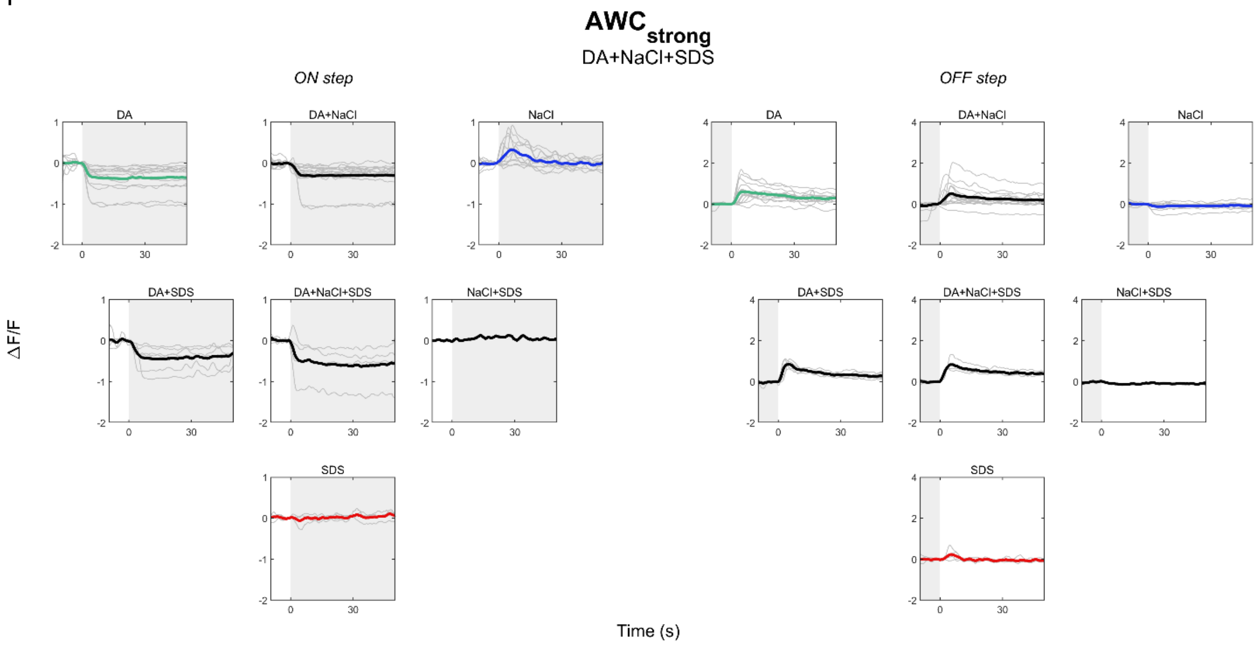


**Supplementary Figure 25.** Raw mean traces of AWC$_{strong}$ in NaCl (blue), DA (green) and SDS (red), and their mixtures (black). Individual neuron traces in grey. Shaded area indicates presence of stimulus.

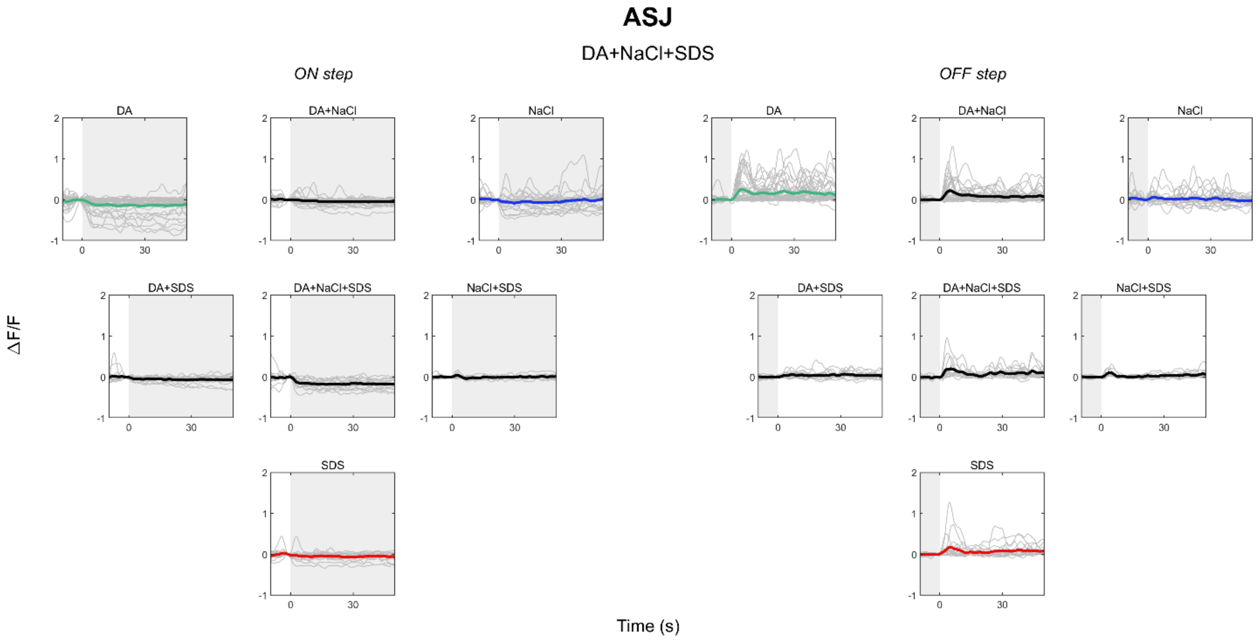


**Supplementary Figure 26.** Raw mean traces of ASJ in NaCl (blue), DA (green) and SDS (red), and their mixtures (black). Individual neuron traces in grey. Shaded area indicates presence of stimulus.

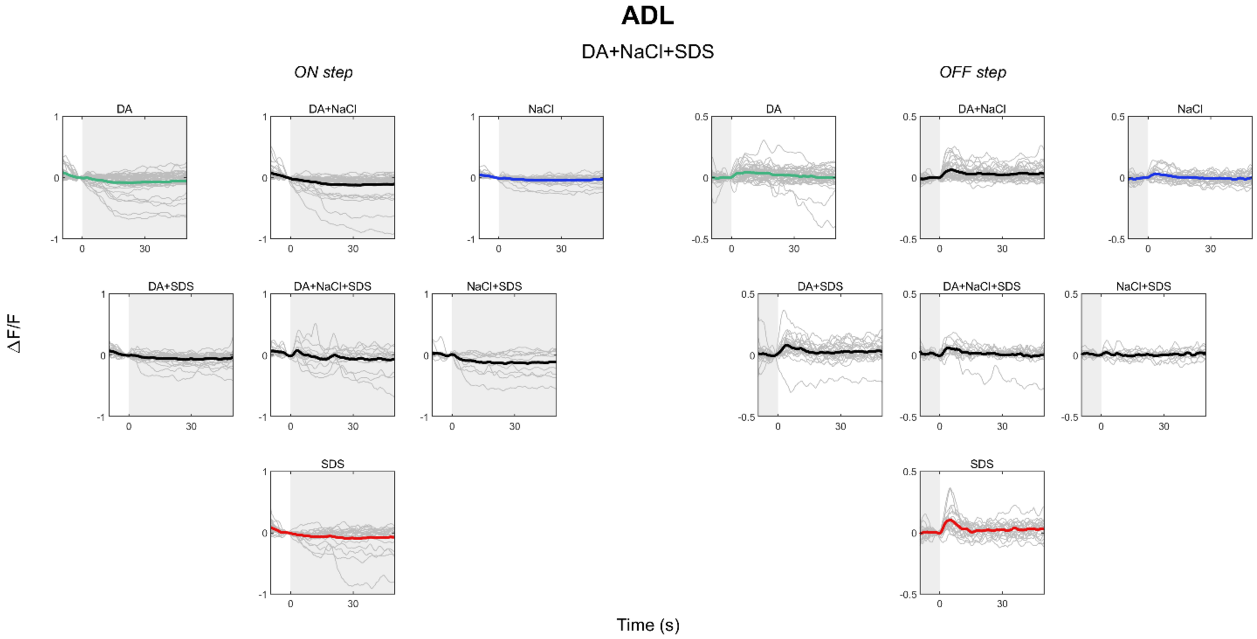


**Supplementary Figure 27.** Raw mean traces of ADL in NaCl (blue), DA (green) and SDS (red), and their mixtures (black). Individual neuron traces in grey. Shaded area indicates presence of stimulus.

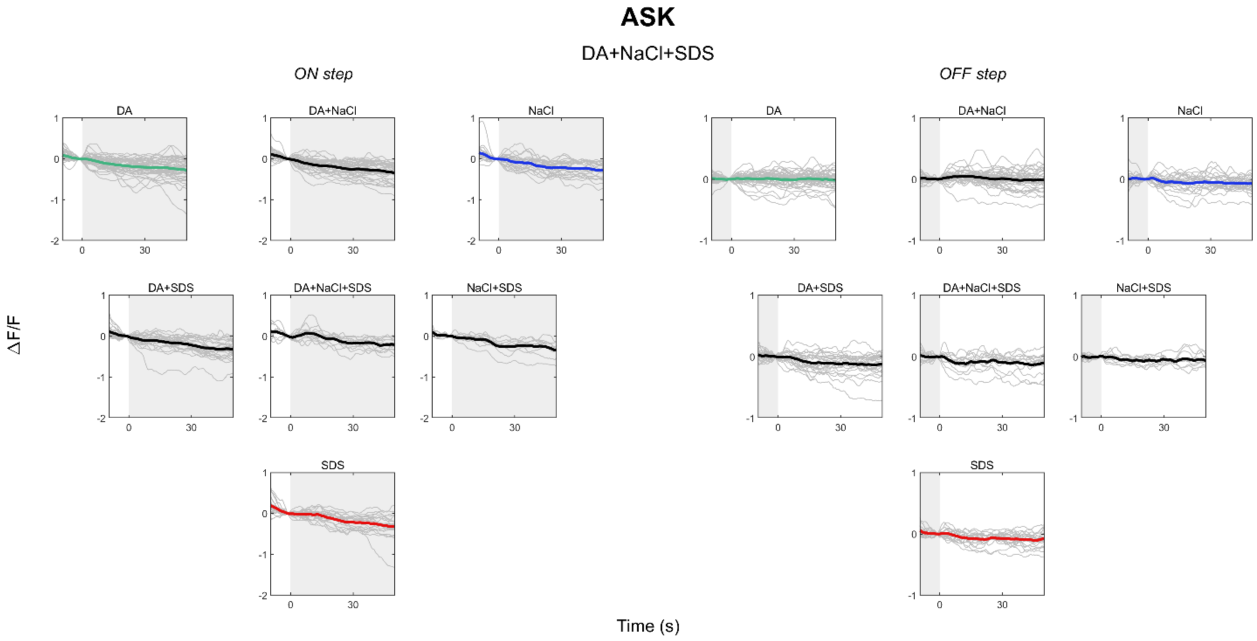


**Supplementary Figure 28.** Raw mean traces of ASK in NaCl (blue), DA (green) and SDS (red), and their mixtures (black). Individual neuron traces in grey. Shaded area indicates presence of stimulus.

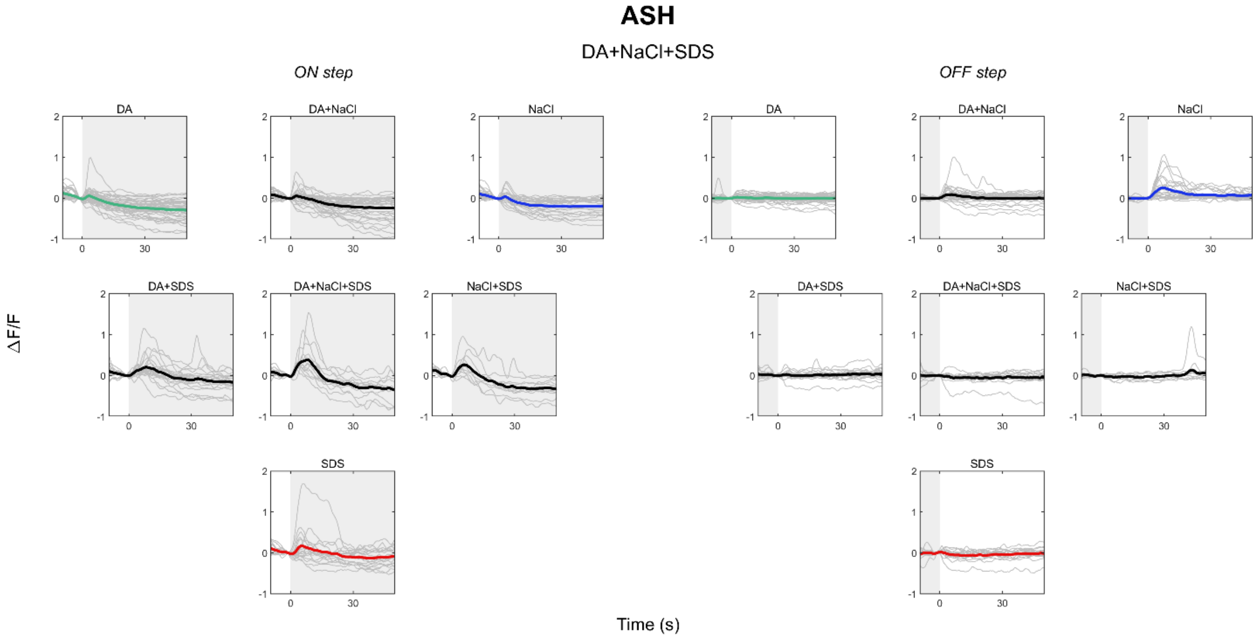


**Supplementary Figure 29.** Raw mean traces of ASH in NaCl (blue), DA (green) and SDS (red), and their mixtures (black). Individual neuron traces in grey. Shaded area indicates presence of stimulus.

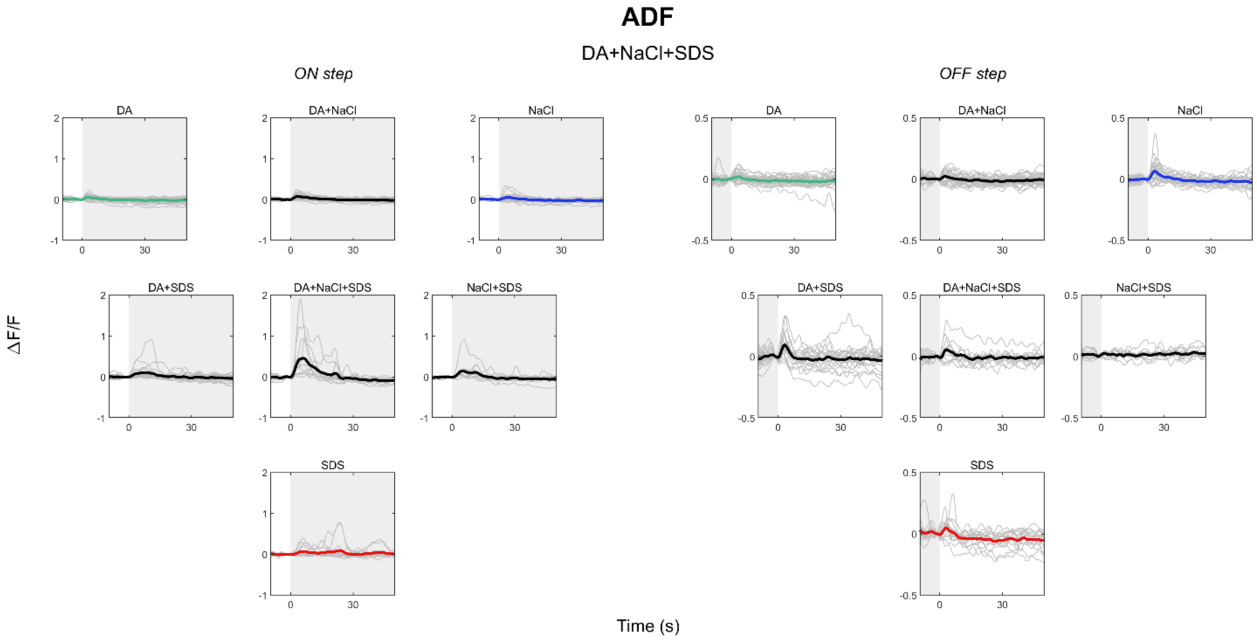


**Supplementary Figure 30.** Raw mean traces of ADF in NaCl (blue), DA (green) and SDS (red), and their mixtures (black). Individual neuron traces in grey. Shaded area indicates presence of stimulus.

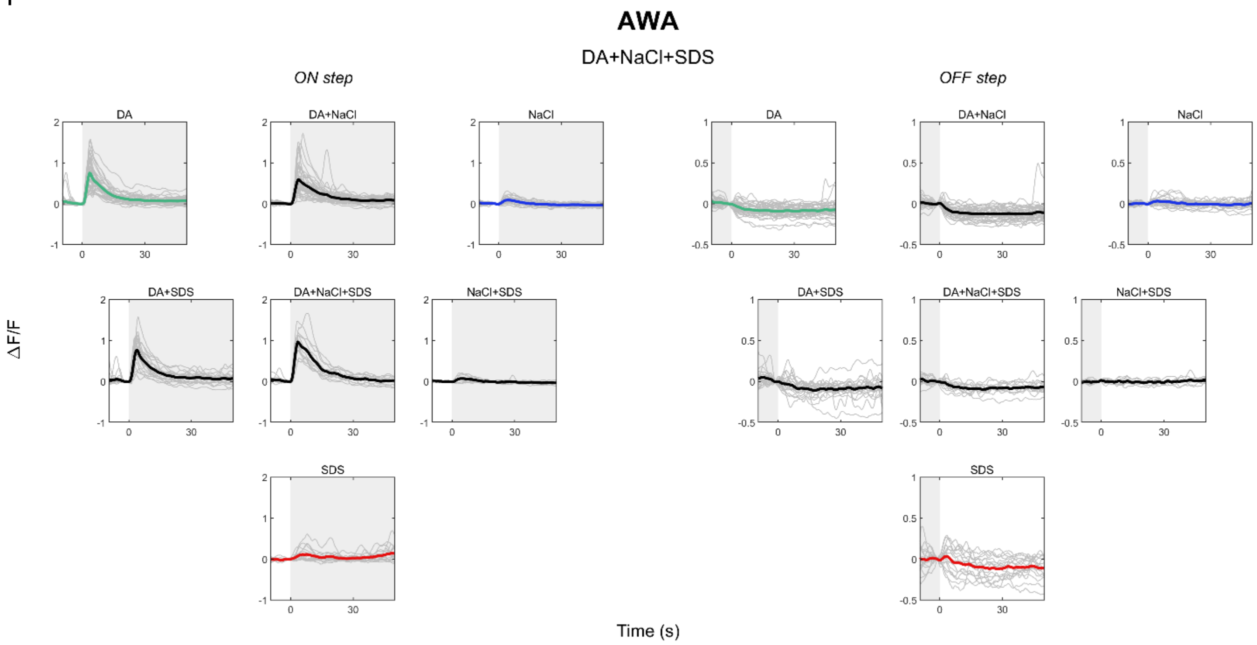


**Supplementary Figure 31.** Raw mean traces of AWA in NaCl (blue), DA (green) and SDS (red), and their mixtures (black). Individual neuron traces in grey. Shaded area indicates presence of stimulus.

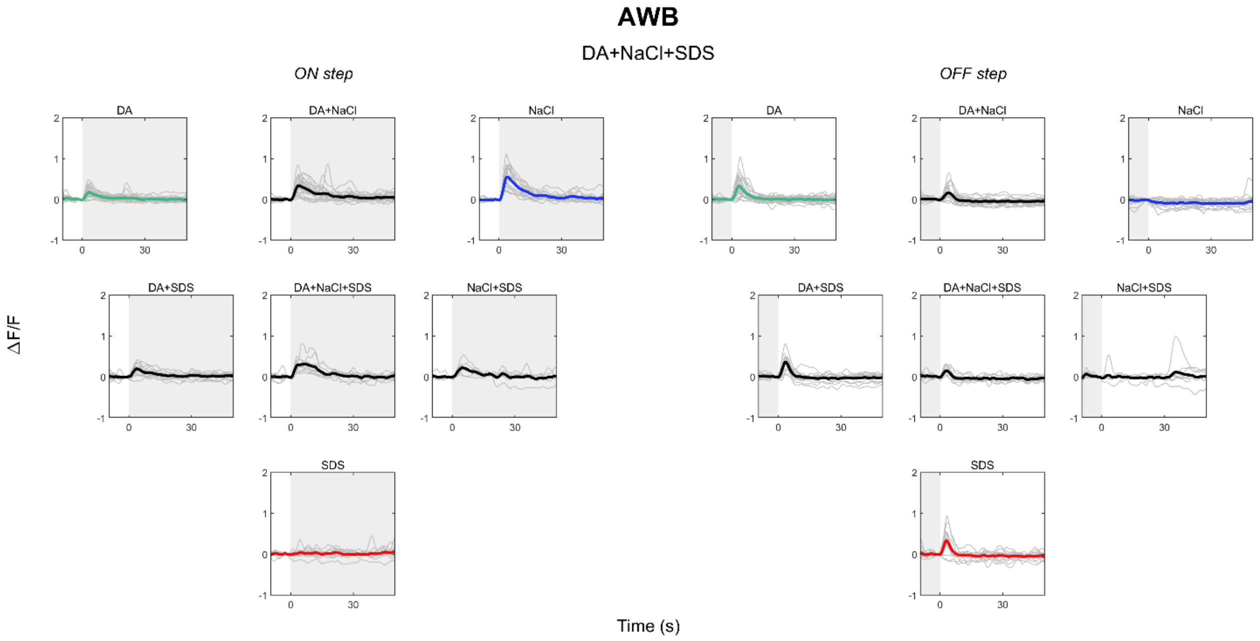


**Supplementary Figure 32.** Raw mean traces of AWB in NaCl (blue), DA (green) and SDS (red), and their mixtures (black). Individual neuron traces in grey. Shaded area indicates presence of stimulus.

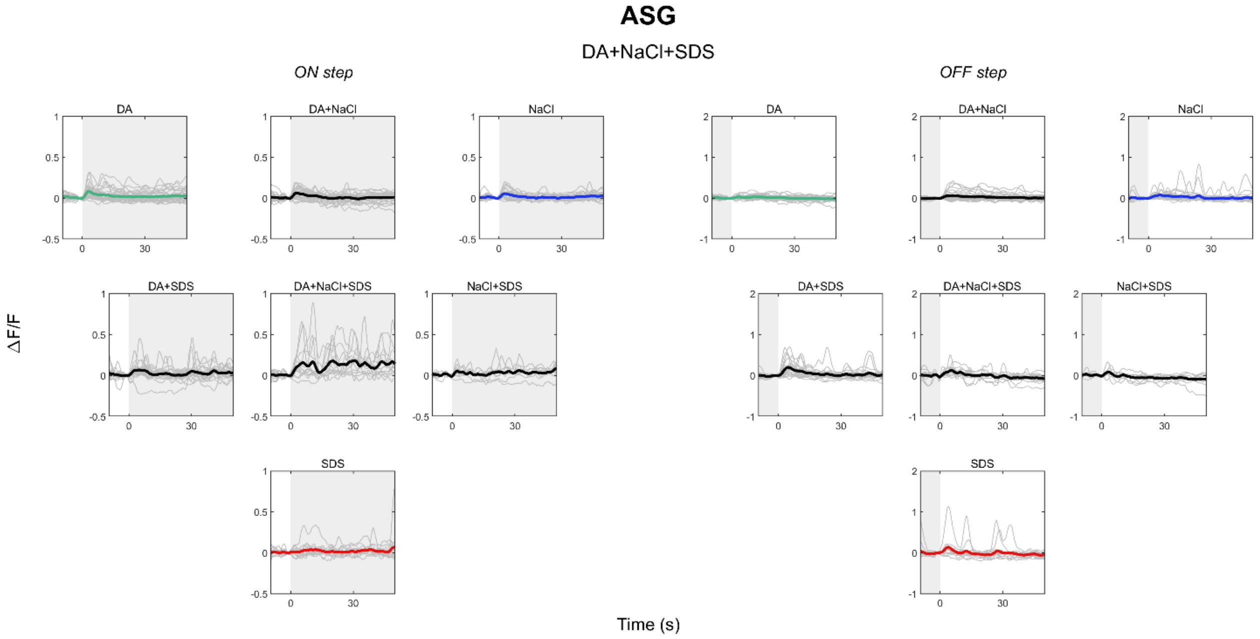


**Supplementary Figure 33.** Raw mean traces of ASG in NaCl (blue), DA (green) and SDS (red), and their mixtures (black). Individual neuron traces in grey. Shaded area indicates presence of stimulus.

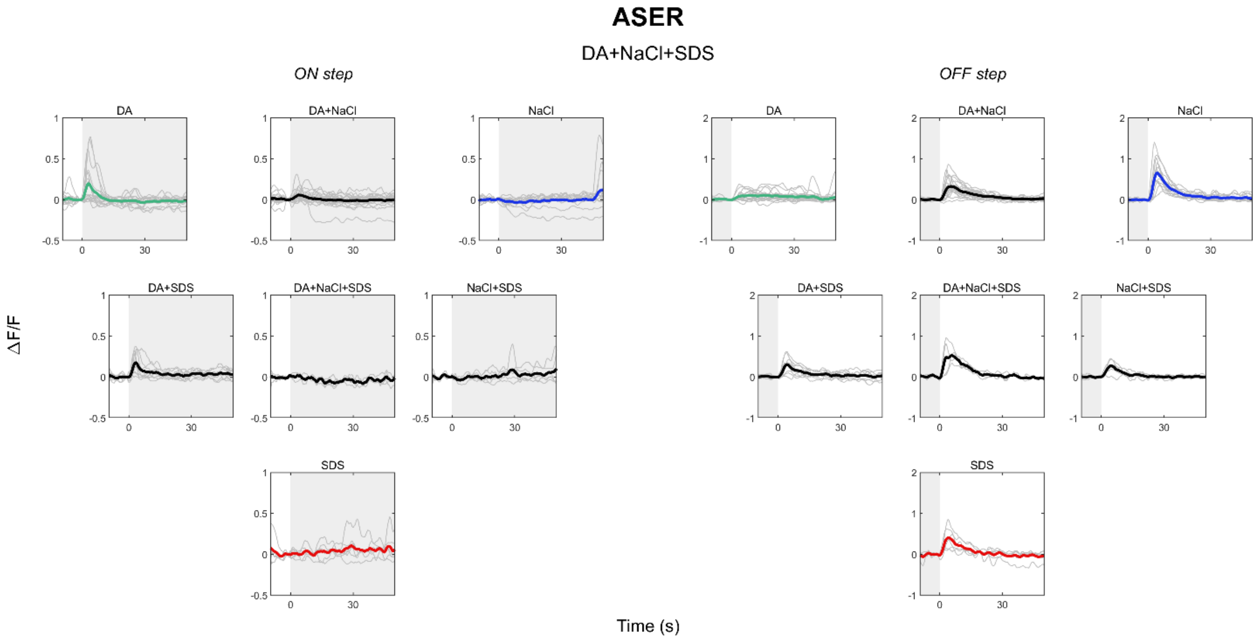


**Supplementary Figure 34.** Raw mean traces of ASER in NaCl (blue), DA (green) and SDS (red), and their mixtures (black). Individual neuron traces in grey. Shaded area indicates presence of stimulus.

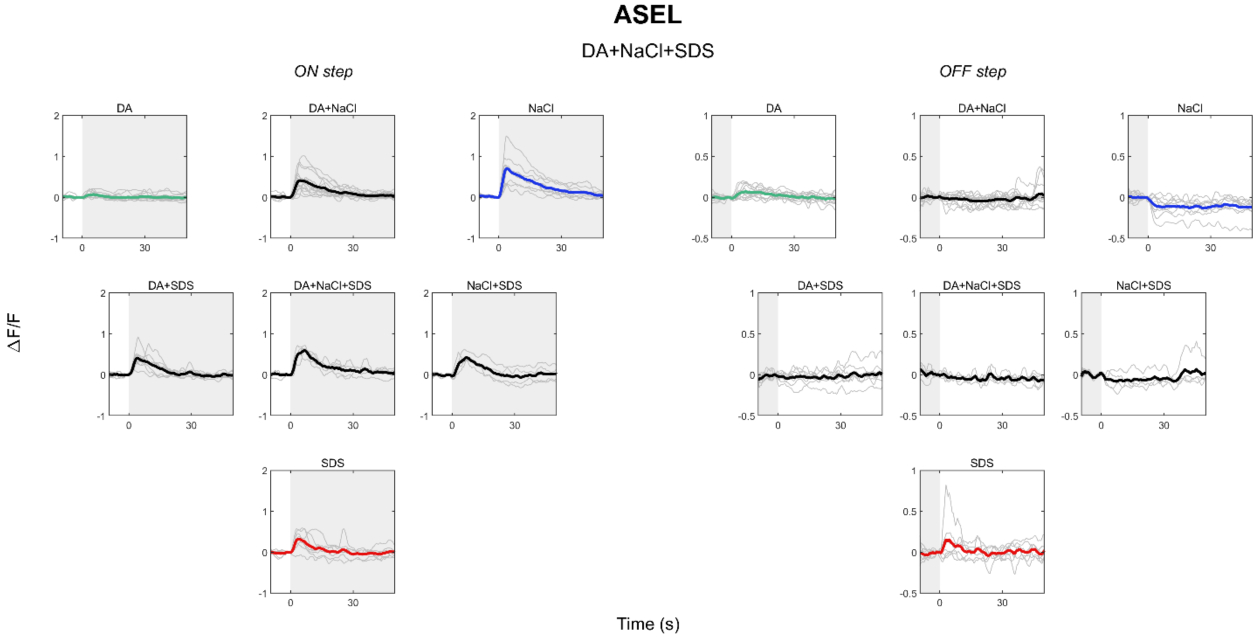


**Supplementary Figure 35.** Raw mean traces of ASEL in NaCl (blue), DA (green) and SDS (red), and their mixtures (black). Individual neuron traces in grey. Shaded area indicates presence of stimulus.

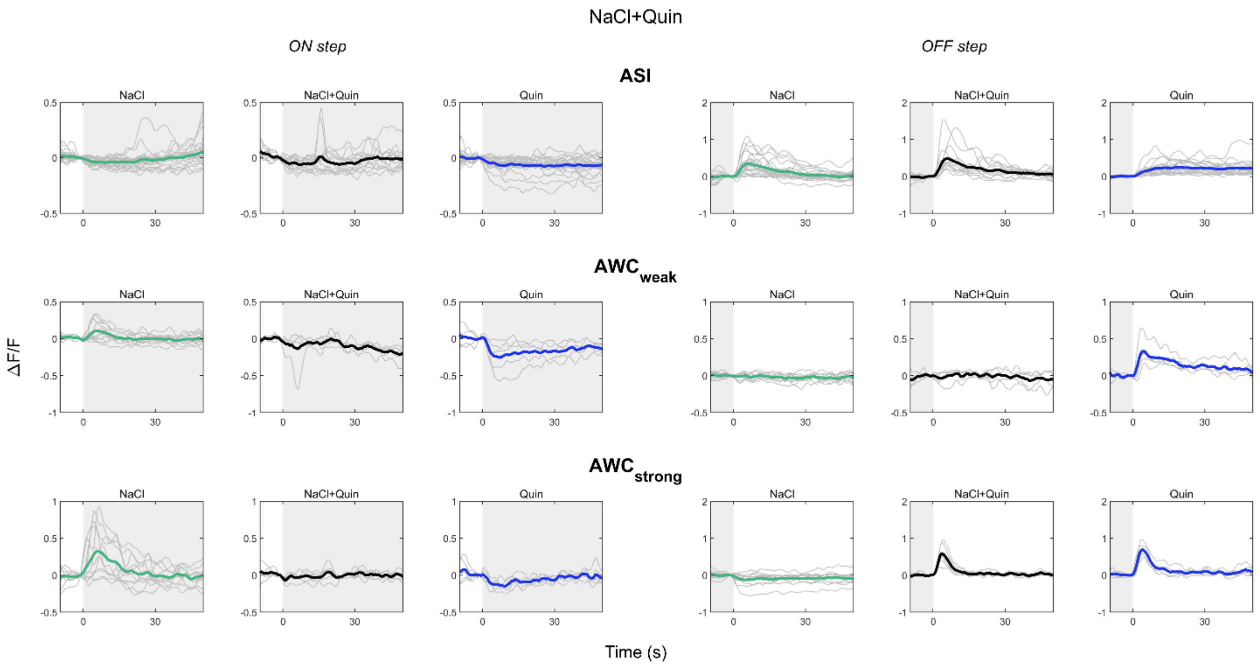


**Supplementary Figure 36.** Raw mean traces of ASI (top), $AWC_{weak}$ (middle) and $AWC_{strong}$ (bottom) in Quin (blue), NaCl (green), and their mixtures (black). Individual neuron traces in grey. Shaded area indicates presence of stimulus.

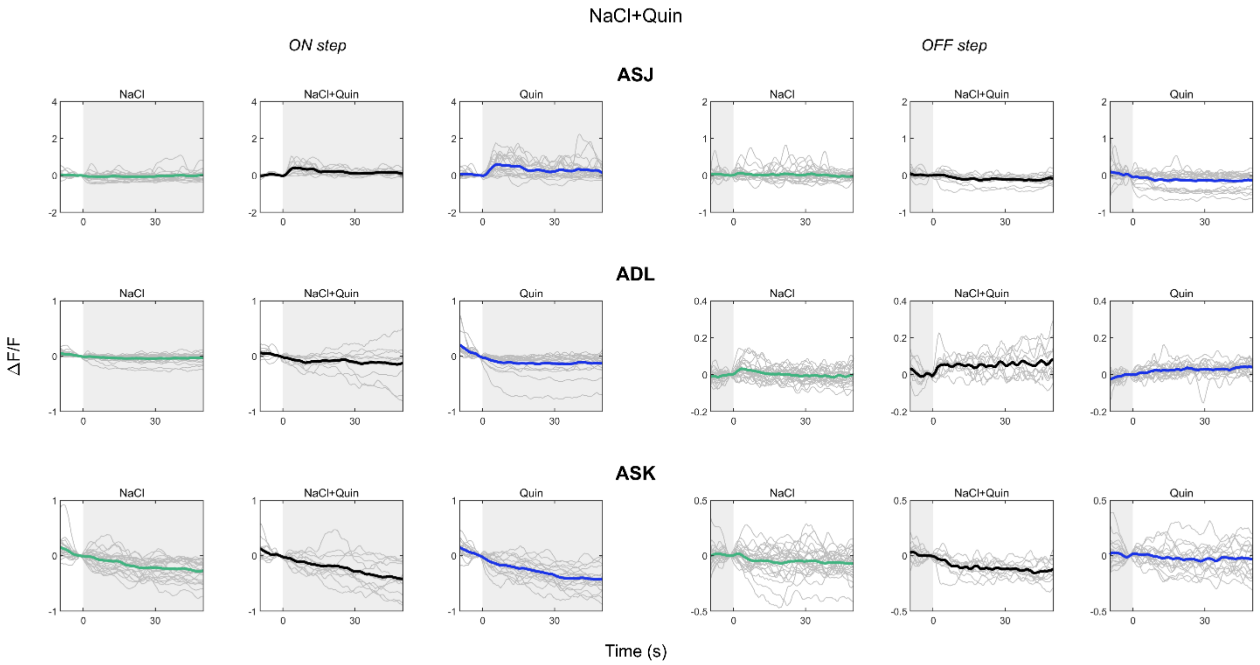


**Supplementary Figure 37.** Raw mean traces of ASJ (top), ADL (middle) and ASK (bottom) in Quin (blue), NaCl (green), and their mixtures (black). Individual neuron traces in grey. Shaded area indicates presence of stimulus.

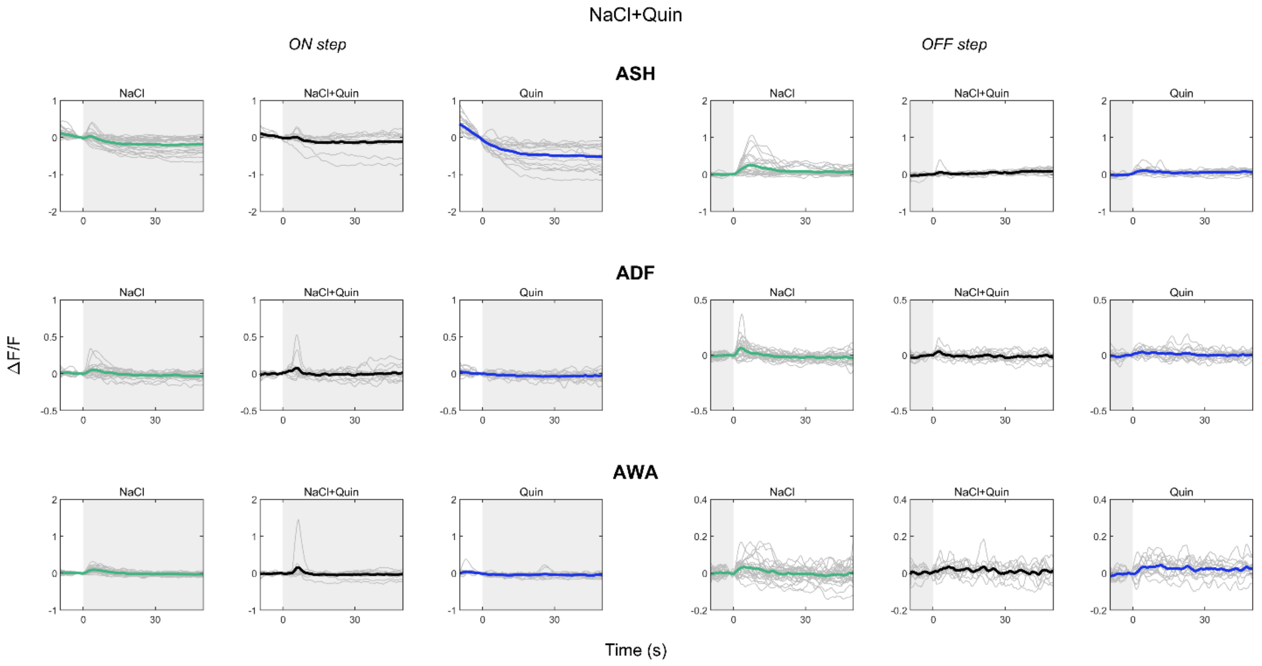


**Supplementary Figure 38.** Raw mean traces of ASH (top), ADF (middle) and AWA (bottom) in Quin (blue), NaCl (green), and their mixtures (black). Individual neuron traces in grey. Shaded area indicates presence of stimulus.

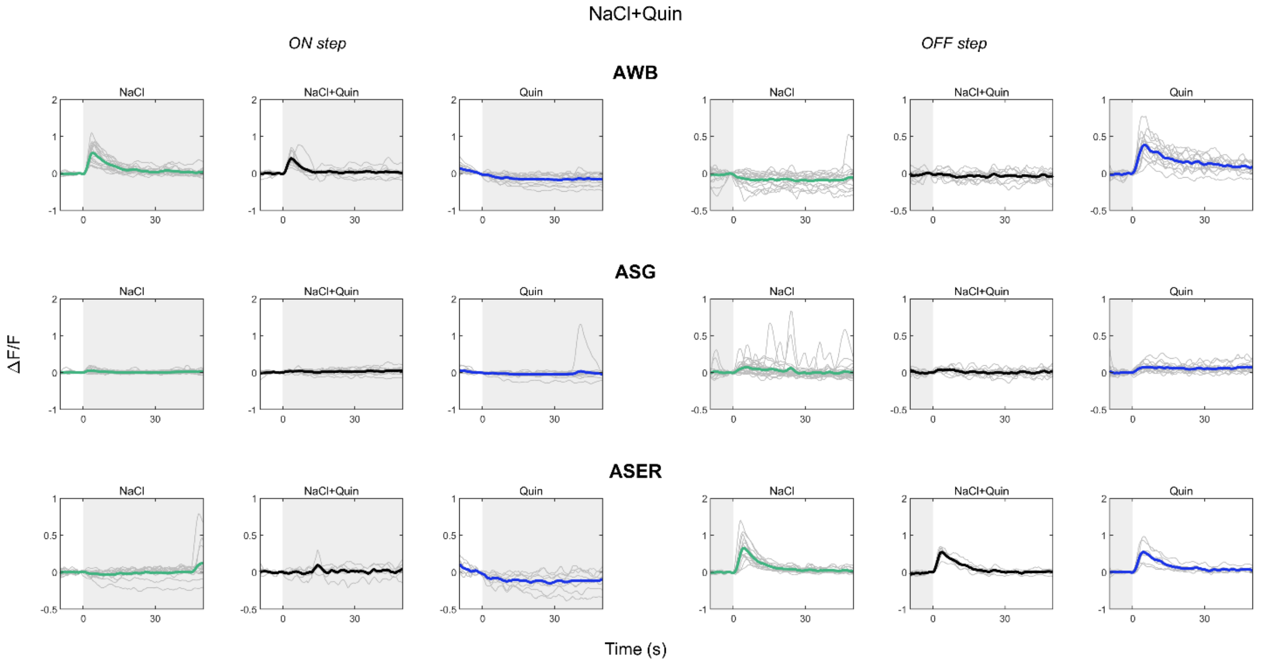


**Supplementary Figure 39.** Raw mean traces of AWB (top), ASG (middle) and ASER (bottom) in Quin (blue), NaCl (green), and their mixtures (black). Individual neuron traces in grey. Shaded area indicates presence of stimulus.

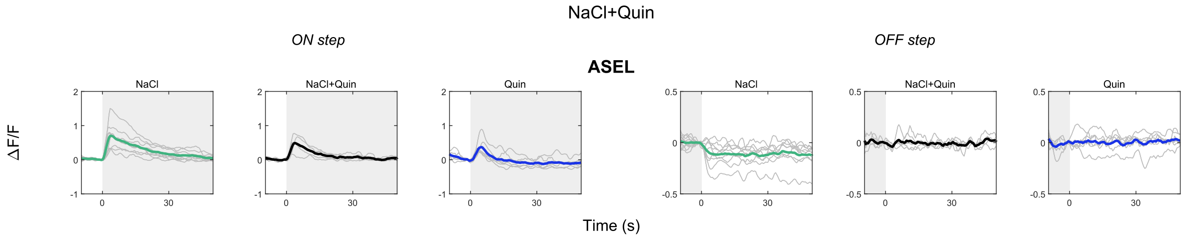


**Supplementary Figure 40.** Raw mean traces ASEL in Quin (blue), NaCl (green), and their mixtures (black). Individual neuron traces in grey. Shaded area indicates presence of stimulus.
